# A primary patient-derived model for investigating functional heterogeneity within the human Leukemic Stem Cell Compartment

**DOI:** 10.1101/2022.03.01.482535

**Authors:** Héléna Boutzen, Michelle Chan-Seng-Yue, Alex Murison, Nathan Mbong, Elvin Wagenblast, Christopher Arlidge, Seyed Ali Madani Tonekaboni, Elias Orouji, Andrea Arruda, Amanda Mitchell, Faiyaz Notta, Mathieu Lupien, Mark D. Minden, Kerstin B. Kaufmann, John E. Dick

**Author notes:** Corresponding Author : John E. Dick, Princess Margaret Cancer Centre, PMCRT, 8th Floor, Rm 8-358, 101 College Street, Toronto, Ontario, M5G 1L7.

## Abstract

The ability of leukemic stem cells (LSC) to evade therapy and fuel leukemic progression causing relapse impedes therapeutic success in acute myeloid leukemia (AML). The LSC pool within a patient sample is not homogenous but comprises distinct LSC subsets that vary in self-renewal and propagation properties. The stemness programs that underlie LSC types are poorly understood since human LSC studies require primary patient samples where LSC numbers are low and isolation methods impure. To overcome these challenges, we developed a patient-derived AML model system (OCI-AML22) displaying a functionally, transcriptionally and epigenetically defined cellular hierarchy driven by functional LSCs that can be immunophenotypically identified and isolated. Through single cell and functional approaches, the OCI-AML22 LSC fraction was found to contain distinct LSCs that vary in proliferative and differentiation properties. OCI-AML22 represents a valuable resource to decipher mechanisms driving stemness and the multiple layers of heterogeneity within LSCs.

## Introduction

Acute myeloid leukemia is a heterogeneous disease (1–6) driven by leukemic stem cells (LSCs) (7–16). Some AML patients can achieve durable remission, but the majority will relapse within 2 years. There is strong evidence that relapse arises from LSCs that are capable of surviving chemotherapy and initiating relapse. (7–16). Consequently, AML survival remains poor and a better understanding of mechanisms fuelling stemness and chemoresistance in LSC is required to design efficient therapies. LSCs from across the spectrum of AML patients can exhibit heterogeneity in frequency and immunophenotype. Moreover, LSCs can be genetically diverse both between patients but also within a single patient where they drive genetic subclones (15). Despite this heterogeneity, all LSCs share the same functional hallmark properties of stem cells: capacity for self-renewal allowing long term generation of disease. These functional properties are clinically relevant because they are directly linked to relapse (15) and poor prognosis (8,9). Many studies from our lab and others have established that the intrinsic stemness properties of LSCs from across many diverse AML cohorts can be captured using gene expression signatures (8,9). Many of these signatures are also shared with normal HSC (8). These LSC signatures are highly prognostic, demonstrating that stemness is capturing a shared property that can be linked to clinical outcome within heterogeneous AML cohorts. The LSC17 score represents the most recent and validated proof of this concept. The score was derived from 236 functionally-defined fractions (138 LSC+ and 98 LSC-fractions) obtained from 78 patients samples. Indeed, this score has been shown to be highly prognostic in multiple independent AML datasets adult or pediatric, altogether spanning >1000 AML samples Ng et al., Nature, 2016: 908 samples; Duployez et al., Leukemia, 2019: 200 samples) (9,17). The LSC17 score remains highly prognostic in multivariate analyses and outperforms previously defined prognostic factors such as age or cytogenetics. Thus regardless of the diverse paths taken during leukemogenesis, the high prognostic power of LSC17 suggests that all leukemogenic pathways converge onto stemness properties. The convergence of intrinsic stemness properties between AML patients has enabled the deployment of the LSC17 score into the clinic (Ng et al., Blood Advances, in press (18).

There has been considerable effort to better understand the key biological processes that distinguish LSC from the other non-LSC leukemic cells including proteostatic responses, epigenetic pathways, immune escape ability or metabolism (5,19–23). A number of these processes have been translated into therapeutic strategies (5,6,24–27). However, significant challenges to study the biology of LSCs remain. First, primary AML cells are difficult to investigate experimentally as they do not expand well in culture. While traditional AML cell lines do not recapitulate the LSC-driven cellular hierarchy of primary AML samples (28). Second, no markers exist to purify rare LSC populations to homogeneity (29). Like many other aspects of AML, the relationship between cell surface phenotype and functionally defined LSC is not consistent across a spectrum of patient samples preventing simple use of immunophenotypic markers as a surrogate for identifying LSC in every patient (29,30). This forces reliance on cumbersome, expensive and time-consuming xenograft assays for LSC detection and quantification by functional analysis. Moreover, we and others have shown that the pool of LSCs is not homogenous. At the genetic level, genetically distinct LSCs co-exist within the same AML patient sample (31–37). At the functional level, LSCs within an individual sample differ in their self-renewal and xenograft repopulation capacity (38). This functional heterogeneity results in the coexistence of distinct types of LSC within each AML sample including a class that is latent, slowly repopulating and another that exhibits faster repopulation kinetics (38). The mechanistic basis for this functional heterogeneity within the pool of LSCs, and the resultant vulnerabilities that might exist for each type of LSC are unknown; knowledge that is central for the design of effective therapies that would eradicate the entire LSC pool (30). Overall, it has been difficult to study LSC functional heterogeneity; they are rare and no purification methods exist to separate functionally distinct types of LSC within the same sample; the requirement to use primary cells and xenograft assays and the inability to propagate LSC *in vitro* have all greatly hindered mechanistic and multi-omic LSC studies. Thus, there is great need for model systems that retain the multiple layers of functional heterogeneity that exist within primary human AML samples.

Here, we report the development of a patient-derived primary AML model, called OCI-AML22, which reflects the functional, transcriptional and epigenetic cellular hierarchy common to most primary AML samples. Importantly, distinct classes of LSC can be identified, opening a powerful resource to uncover the molecular basis for complexity driving the distinct LSC subsets that collectively comprise the LSC pool. OCI-AML22 can also be efficiently modified with lentivectors and CRISPR based methods yielding a novel model for screening and the development of more efficient LSC-targeted therapies.

## Results

### OCI-AML22 models the phenotypic and functional hierarchy found in primary AML

To establish an AML model that recapitulates a cellular hierarchy with LSCs at the apex, we screened AML patient samples (n=34) for their ability to grow and expand in culture (Supplementary Figure S1A and Table 1). Most AML samples could not be maintained for more than a few weeks and no sample consistently expanded in culture (Supplementary Figure S1A), which is aligned with previous observations (28). However, we identified one sample, from a relapse patient who underwent multiple cycles of chemotherapy (see Material and Methods) that could be expanded long term *in vitro* (Supplementary Figure S1A-B). *In vitro* expansion of this sample resulted in an immunophenotypic hierarchy of four fractions defined by CD34 and CD38 cell surface expression (Figure1A and Supplementary Figure S1B). The OCI-AML22 CD34+CD38- fraction was the only fraction able to maintain the entire hierarchy in culture reestablishment assays (Figure 1B-C).

**Figure 1:**
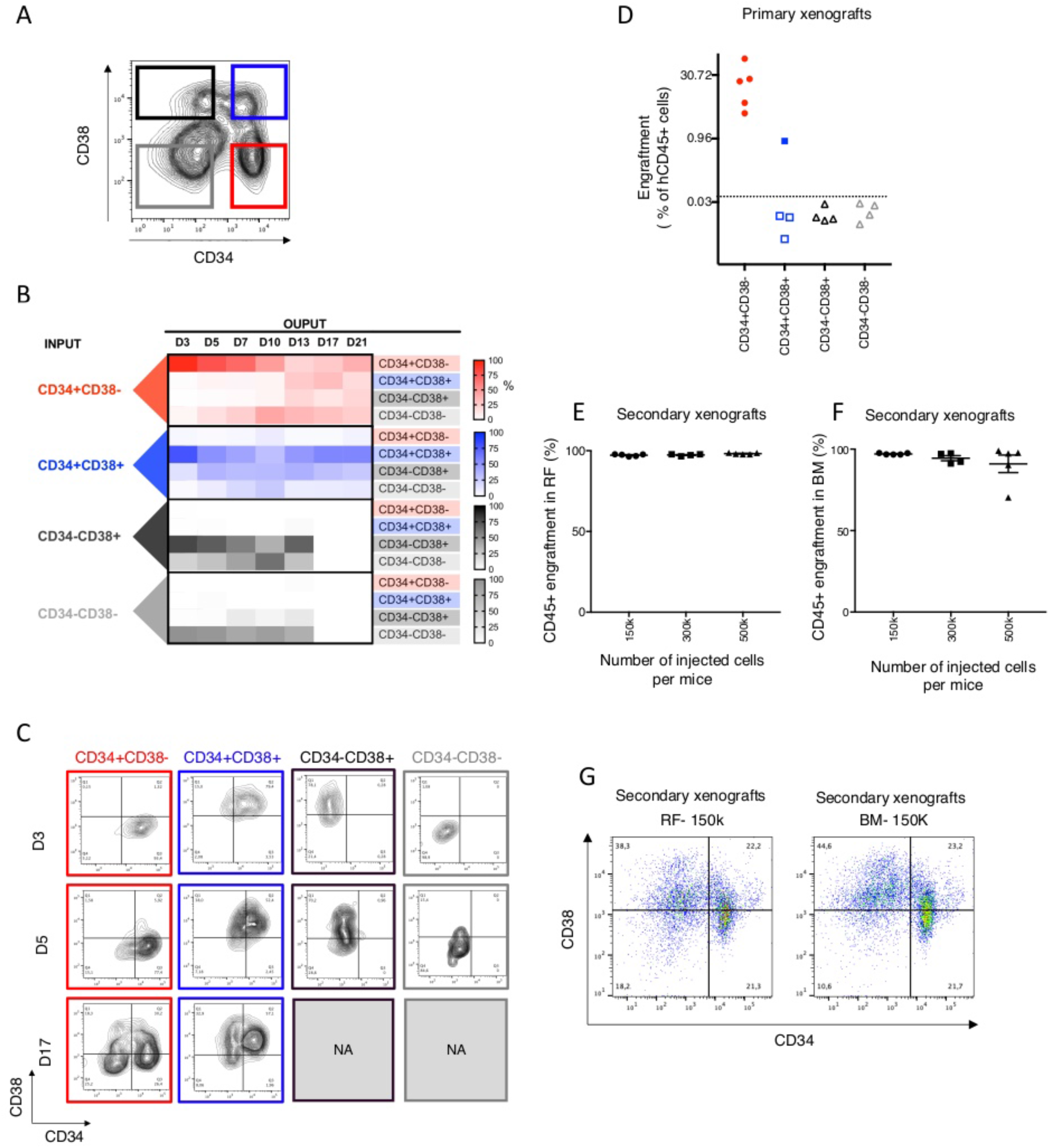
OCI-AML22 models the phenotypic and functional hierarchy found in primary AML. **A**. Immunophenotypic profile of OCI-AML22 with the 4 fractions sorted as indicated based on CD34 and CD38 cell surface expression. **B.** Immunophenotypic characterization of cellular outputs over time generated by each sorted fraction following in vitro expansion as assessed by flow cytometry. **C.** Flow cytometric profiles of sorted fractions over time; related to (B). *NA* = *not available due to no viable cells remaining*. **D.** NSG mice were injected with sorted populations (100 000 cells per mouse) as indicated. Engraftment level was assessed 8 weeks after injection by flow cytometry measuring the percentage of human CD45+ (hCD45) cells in the injected femur. Each dot on the graph represents a mouse. **E-G.** Cells collected from the xenografts generated after injection of OCI-AML22 fractions from Fig 1D were pooled and sorted for human cells then injected into NSG-SGM3 mice at the indicated cell dose per mouse. Engraftment level (AnnexinV-, 7AAD-, hCD45+) was assessed 8 weeks later in the injected bone (RF) (E), non-injected bone, referred to as bone marrow (BM) (F). Representatif FACS profiles of grafts in the right femur (left) or the bone marrow (right) are represented. (G).

To determine whether the immuno-phenotypic hierarchy also reflected a functional LSC-driven hierarchy, the four CD34/CD38 subpopulations were sorted after 3-4 months of *ex vivo* expansion and assessed for their leukemia-initiating capacity (L-IC) using xenotransplantation assays (Figure 1D). All NSG mice (5/5) injected with 100,000 CD34+CD38- cells were engrafted at 8 weeks. By contrast, injection of the same number of CD34+CD38+ or CD34- cells resulted either only in a single engrafted mouse (1 of 4 injected mice, 0.8% engraftment) or in no detectable engraftment, respectively (Figure 1D). Serial transplantation of cells harvested from primary mice resulted in robust engraftment of secondary recipients (Figure 1E-F) indicating that L-IC had self-renewal potential. The prospective purification of self-renewing L-IC formally establishes their identity as LSC (39) and confirms that this AML model (named OCI-AML22) is organized as a functional cellular hierarchy driven from LSC at the apex. The LSCs can be easily phenotypically identified and purified by sorting CD34+CD38- cells. Differentiation profiles of secondary grafts also establish the capacity to recapitulate the entire hierarchy *in vivo* (Figure 1G). To quantify the absolute LSC content within the CD34+CD38- fraction of OCI-AML22, we determined LSC frequency using a limiting dilution xenograft assay (Table 2). The average LSC frequency was estimated to be 1/286 (range: 1/102-1/804) 12 weeks after injection. When compared to LSC frequencies obtained for 45 LSC containing fractions of primary AML samples (9,40), the OCI-AML-22 LSC fraction is at the high end of the spectrum of LSC potential (Supplementary Figure S2A). Collectively, these data establish that OCI-AML22 exhibits a phenotypic and functional hierarchy and has the capacity for both *in vitro* and *in vivo* long-term propagation.

### OCI-AML22 maintains the polyclonal genetic architecture of the primary AML sample

Since the donor sample from which OCI-AML22 was derived had complex cytogenetics, we assessed the genetic stability of the OCI-AML22 model during *in vitro* and *in vivo* expansion and compared the subclonal genetic structure of OCI-AML22 with the donor. Whole genome sequencing (WGS) was undertaken on the primary donor sample (day 0), the cultured OCI-AML22 model (~day 30), as well as on xenografts generated from CD34+CD38- cells sorted from the cultured sample (~d120) (Figure 2A). Genomic analysis showed that cultured and donor samples were highly similar with 99% conservation. The CD34+ fraction from xenografts showed an even higher conservation (99.8%). Detailed analysis further revealed that the minor genetic differences between the dominant clones of the cultured, xenografted and donor samples were reflective of subclonal genetic diversity where the dominant populations of the culture and xenografts arose from rare pre-existing subclones present in the donor sample (Figure 2B-D, Supplementary Figure S3A-B and Table 3-4). There is precedence for this finding from our studies of B-ALL xenografts (41). WGS showed that the dominant clone in the OCI-AML22 cultured sample exhibits a series of amplifications on chromosome 11 (Supplementary Figure 3, arrow 1). This clone was termed clone AMP11 (Supplementary Figure S3, line 2 and Figure 2B, line2, in red). This series of amplifications on chromosome 11 are not present in the dominant clone of the donor sample (Supplementary Figure S3, arrow 1, line 1). However, in depth analysis of this region on chromosome 11 in the donor sample showed that these alterations were already present at a subclonal level (Figure 2C and Table 3-4). Thus the series of 11q amplifications were not generated by a genetic drift due to culture, but arose by the selective amplification of a minor clone preexisting in the donor sample. Of note, the AMP11 clone could also be detected at the subclonal level in xenografts generated by the cultured sample (Table 3-4). WGS also revealed that the dominant clone in the donor (Supplementary Figure S3A, arrow 1) contains a small deletion on chromosome 5 (and lacks the 11q amplification) (Supplementary Figure S3A, arrow 2); termed clone del5 (Supplementary Figure S3, line1 and Figure 2B, line 1, in blue). By contrast, the dominant clone in xenografts, generated by the cultured sample, lacked both the chromosome 5 deletion and the chromosome 11q amplifications (Supplementary Figure S3A, line 3). Since a deletion cannot be spontaneously restored to normal, the clone that preferentially expands *in vivo* must be ancestral to both the del 5 and the AMP11 clones. This clone is termed the ancestral clone (Supplementary Figure S3A, line 3 and Figure 2B, line3 in yellow). Thus, *in vitro* culture, followed by *in vivo* propagation of the donor sample, provided insight into the evolutionary relationships of the three genetic subclones present at different levels in the original donor sample (Figure 2D). These include the del5 clone, dominant in the donor sample (Figure 2B and 2D-E, in blue), the minor AMP11 clones, and the ancestral clone, that can generate both del5 and AMP11 subclones. (Figure 2B and 2D-E, in yellow). The ancestral clone must be present in both the donor and in culture at subclonal levels since it becomes detectable following xenotransplantation of the cultured OCI-AML22 cells (Figure 2E, in yellow). The AMP11 clone, present at the subclonal level in the donor, preferentially expands in culture and remains present but at the subclonal levels in xenografts generated by the cultured sample (Figure 2B and 2D-E, in red). Altogether, the WGS analysis revealed that the cultured OCI-AML22 model is genetically stable during prolonged *in vitro* and *in vivo* propagation while maintaining at least 2 of the clones originally present in the donor sample: the ancestral clone and the AMP11q clone. These results are reminiscent of our findings documenting leukemia evolution using xenografts (41) and highlight that the OCI-AML22 model maintains the polyclonal genetic architecture characteristic of primary AML samples in contrast with traditional AML cell lines that are clonal.

**Figure 2:**
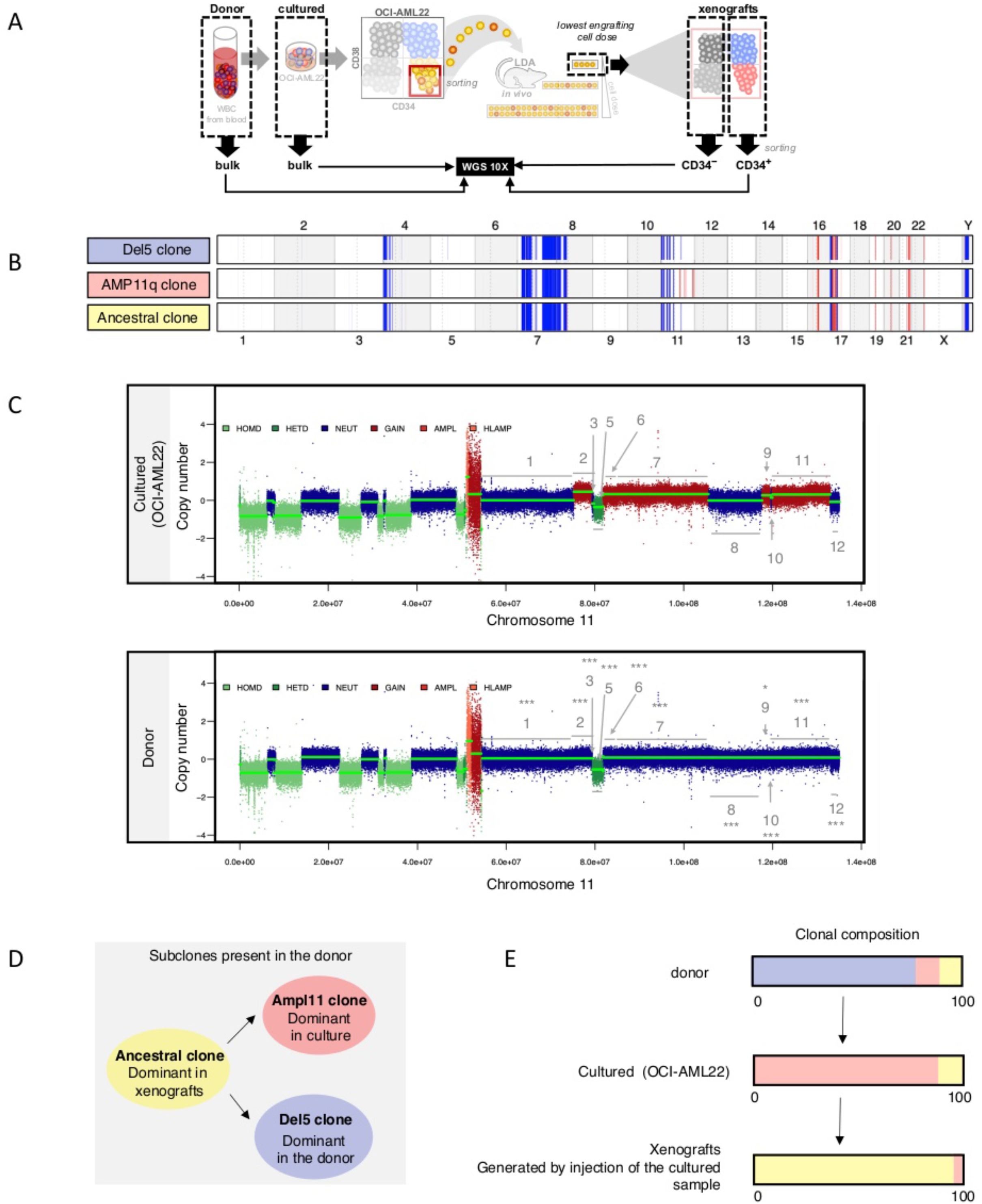
OCI-AML22 maintains the polyclonal genetic architecture of the primary AML sample. **A.** Schematic representation of the samples sequenced for WGS. **B**. Alterations present in each of the dominant clones are displayed. Copy number losses are defined as CN less than 1.5 (shown in blue) while gains have CN greater than 2.5 (shown in red). The width of the colored region corresponds to the size of the modified region in the genome. **C.** Representation of the copy number obtained on chromosome 11 for cultured and donor samples. Segment regions that were detected using hmmcopy in the cultured sample (top) were used to break down the chr11 arm q of the donor sample (bottom) into matching regions. Within these segments, each dot (at 1kb intervals) was taken to compare the average copy number to the adjacent segments and thus determine if the average copy numbers were different between the indicated adjacent regions. wilcox tests were run. Each start on top of each region shows the existence of amplifications that can be significantly detected at a subclonal level in the donor sample. **D.** Evolutionary relationships of subclones present in the donor sample. The sample where the clone is dominant is indicated under. **E.** Clonal composition of the different sequenced sample using the same clones color coding as in B and D.

### The OCI-AML22 CD34+CD38- fraction preserves the transcriptional and epigenetic landscape of primary stem cells throughout ex vivo expansion

To determine if the transcriptional signature of the CD34+CD38- fraction was maintained over time, we performed deep RNA sequencing (RNA-seq) of CD34+CD38- cells from the donor sample and of four immunophenotypic fractions harvested at multiple time points of culture (12, 60, 90 days) (Fig 3A). Principal component analysis (PCA) showed that even after 2 to 3 months of *in vitro* culture, all CD34+CD38- fractions clustered together with CD34+CD38- cells from the donor sample indicating that *ex vivo* expanded CD34+CD38- cells preserved the global transcriptomic landscape of its original donor (Fig 3B). In accordance with our functional data (Figure 1), CD34+CD38+ populations were positioned in the PCA space between the CD34+CD38- engrafting and non-engrafting CD34- fractions. CD34- fractions clustered together and were the most distinct from CD34+CD38- populations. The functional hierarchical organization described in Figure 1 prompted us to investigate whether OCI-AML22 recapitulates LSC and non-LSC features extracted from heterogenous primary AML cohorts, as well as the diverse AML cellular states recently extracted from scRNA-seq of primary AML samples (10). Gene set variation analysis (GSVA) on each of the OCI-AML22 fractions showed that the OCI-AML22 model recapitulates the various cellular states extracted from primary AML samples with enrichment of primitive states in the functional LSC fractions (HSC, Progenitor, summarized in HSC.Prog) and progressive enrichment for mature states in the LSC downstream progeny (GMP, Promono, Monocyte, summarized in Myeloid) (Fig 3C-D and Supplementary Figure S4A-E). Similarly, using chromatin accessibility signatures generated from highly purified fractions obtained from the normal hematopoietic system (42), we show that the OCI-AML22 CD34+ fraction was enriched in stem cell signatures (LT/HSPC and Act/HSPC), while the OCI-AML22 CD34- non-engrafting fraction was enriched for signatures of mature populations of the myeloid branch (granulocyte, monocyte signatures) (Supplementary Figure S4F). All OCI-AML22 cultured fractions showed the least concordance with the erythroid and lymphocyte (T cells and B cells) signatures compared to other mature populations that belong to the myeloid pathway (granulocyte, monocyte). This result is concordant with a block of differentiation and a shift towards myeloid pathways characteristic for AML (Supplementary Figure S4F).

**Figure 3:**
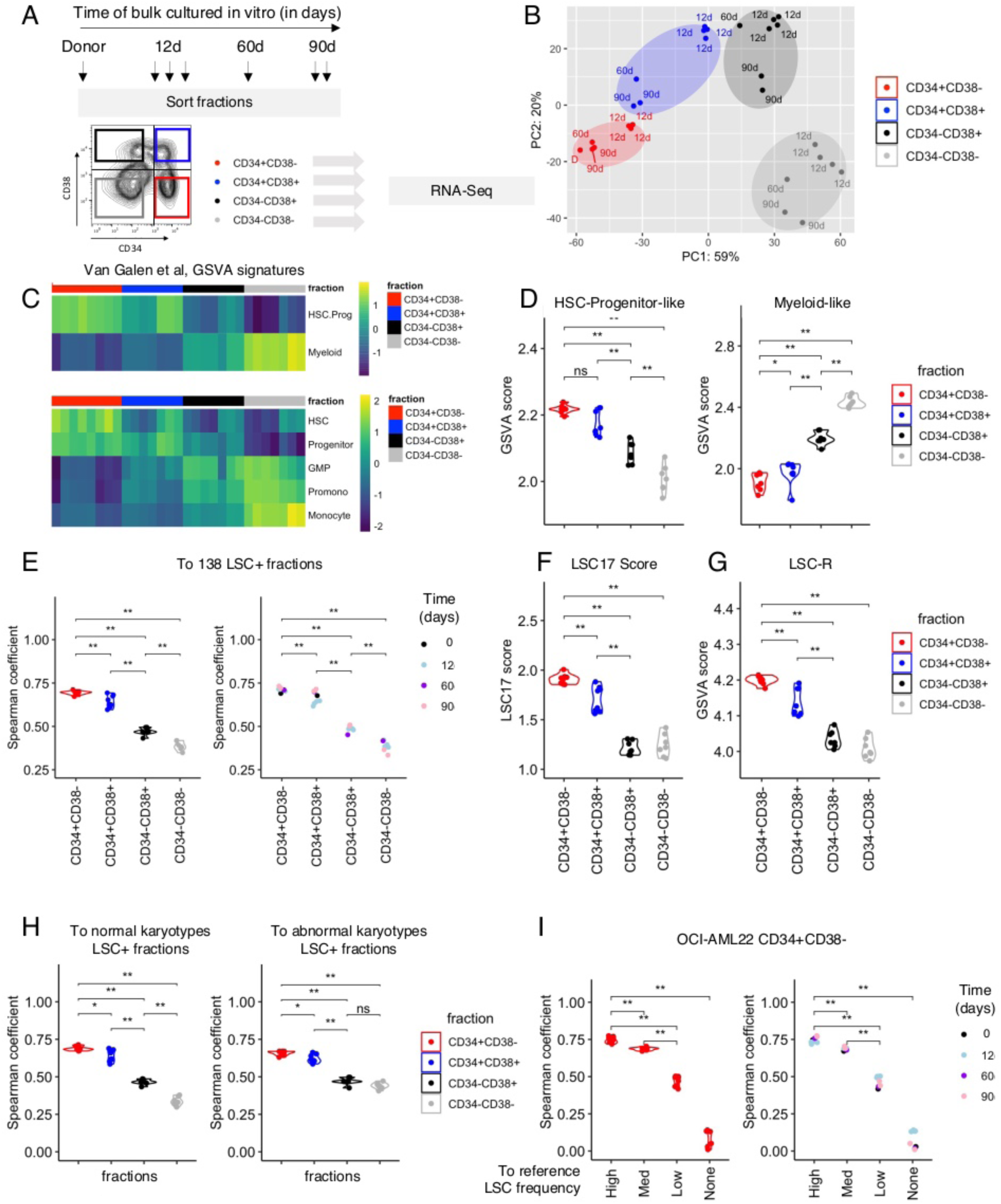
The OCI-AML22 CD34+CD38- fraction preserves the transcriptional and epigenetic landscape of primary stem cells throughout ex vivo expansion. **A.** Schematic representation of OCI-AML22 sorting strategy applied for RNA-Seq. Each arrow indicates an independently expanded culture. These fractions are used throughout the RNA-Seq analysis. **B.** Principal component analysis (PCA) of RNA-seq data generated from CD34+CD38- (red), CD34+CD38+ (blue), CD34-CD38+ (black) and CD34-CD38- (grey) subpopulations. **C.** Supervised heatmap clustering for GSVA scores calculated for the *van Galen* signatures (10) and organized based on OCI-AML22 sorted fraction. Pathways indicative of primitive-like AML cells (HSC-like, Progenitor-like are combined into the HSC-Prog-like signature) while the others (GMP, Promo and Monocyte) are combined in the Myeloid signature. **D-E.** GSVA score of genes present in the HSC-Progenitor-like (D), Myeloid-like (E) signatures (Van galen et al) across OCI-AML22 fractions. **F-G.** Spearman correlation calculated for the gene expression of the LSC104 genes (9) of each OCI-AML22 fractions compared to the average gene expression of these genes across 138 primary AML fractions, enriched for functional LSC described in (9). Fractions are colored depending of the fraction type (F) or the time OCI-AML22 has been maintained in culture before the sort (G). **H**. LSC17 score calculated for each OCI-AML22 fraction. **I**. Gene set variation analysis (GSVA) score of genes present in the LSC-R signature (8) across OCI-AML22 fractions. **J-K.** Spearman correlation coefficient calculated between the LSC104 signature of each of the OCI-AML22 indicated fraction at the bottom, and the LSC104 signature from a group of normal karyotype LSC+ (60 fractions) (J) or a group of abnormal karyotype LSC+ fractions (55 fractions) (K). Normal karyotype LSC+ signatures and abnormal karyotypes LSC signatures are detailed in the methods section. **L-M.** Spearman correlation coefficient comparing the LSC104 signature of the OCI-AML22 CD34+CD38− fractions, to each of the LSC104 signatures for groups of samples indicated on the bottom (LSC frequency High, Medium (Med), low or no detectable LSC activity (LSC neg). Points are colored according to the OCI-AML22 fraction sorted (L) or depending on the time OCI-AML22 has been kept in culture before sorting the CD34+CD38− fraction; 12d: 12 days, 60d: 60 days, 90d: 90 days, P: primary donor)(M).

To determine whether OCI-AML22 CD34+CD38- fractions recapitulate the stem cell transcriptional programs shared by the LSCs from a diverse cohort of primary AML samples, we used the LSC104 stem cell signature. This signature was derived from the largest panel of functionally assessed primary AML fractions currently available in the literature. Additionally, its power to extract the intrinsic transcriptional determinants of stem cells shared across a broad range of functional LSCs and that are clinically relevant has been demonstrated in multiple cohorts (9), making it the ideal tool to determine the broad applicability of the OCI-AML22 model. We first generated an LSC+ signature reference from primary samples by calculating the average expression for each gene in the LSC104 signature across all the 138 LSC+ fractions from Ng et al., Nature, 2016 paper (9). In parallel, using the same approach, we generated LSC104 signatures for each of the OCI-AML22 fractions. We then determined the Spearman correlation coefficient between the primary LSC+ reference group and each of the OCI-AML22 fractions. The resulting correlative score was the highest when comparing the OCI-AML22 CD34+CD38- fractions to the LSC+ reference; the score gradually decreased alongside the reduced engraftment ability of the fractions with the lowest coming from the the non-engrafting CD34-CD38- fractions (Figure 3E, left). These results remained consistent, independent of the time the OCI-AML22 model was maintained in culture. (Figure 3E, right). We reached the same conclusion using 3 independent approaches: LSC17 scoring across all fractions (Figure 3F); GSVA scoring using the LSC–R signature, a previous stemness signature (Figure 3G) (8); and GSVA scoring using the HSC-R signature, a stemness signature generated from normal hematopoietic stem cells (8)(Supplementary Figure S4G).

As OCI-AML22 originated from a genetically complex relapse sample, to demonstrate that the OCI-AML22 LSC fraction is not only a good model of LSCs obtained from complex cytogenetic AML patients, but can also capture the stemness component extracted from AMLs with different karyotypes, we compared the LSC104 signature of the CD34+CD38- OCI-AML22 fraction to the LSC104 signature generated from primary LSC+ fractions obtained from normal karyotype (60 fractions) or abnormal karyotype AMLs (55 fractions) (Figure 3H) where both groups presented similar LSC frequencies (Supplementary Figure S5A). The OCI-AML22 LSC fraction captures the stemness components of primary AML samples, regardless of their cytogenetic status (Figure 3H).

Finally, since the LSC+ fractions from Ng et al (9) that we have used in our study present a broad range of LSC frequencies, we determined the power of the OCI-AML22 LSC fraction to capture the intrinsic transcriptional profile of the most purified LSCs. We generated LSC104 signatures for 4 groups distributed on the basis of LSC frequency: none, low, medium, and high. The LSC104 transcriptional profile of the highest LSC frequency group correlated the best with the OCI-AML22 CD34+CD38- fraction (Figure 3I, left), and this was independent of the time OCI-AML22 cells were maintained in culture (Figure 3I, right). It highlights that the OCI-AML22 LSC fraction captures the transcriptional programs of highly purified LSCs fractions and that these intrinsic stemness components are preserved despite culture. The only criteria that was affecting how well the OCI-AML22 LSC fraction compares to primary AML fractions was LSC frequency, since the primary LSC+ fractions used in our analysis exhibit a wide diversity of clinical and cytogenetic properties (Supplementary Figure S6). Collectively, the OCI-AML22 LSC fraction recapitulates both the transcriptional and epigenetic stemness profile of stem cells extracted from a broad range of primary samples.

### Efficient genetic modification of OCI-AML22 for functional interrogation of leukemia stemness

To determine if we can use the OCI-AML22 model to interrogate molecular determinants driving stemness, we tested a variety of biological properties that were previously reported to discriminate populations within the cellular hierarchy of primary AML samples. We first tested the potential of the OCI-AML22 model to give insights into LSC-specific immune evasion properties since recent studies of 177 primary AML samples showed that LSC, but not their nonstem cell progeny, are able to evade Natural Killer (NK)-driven anti-tumor immunity. LSCs upregulate PARP1 whose overexpression downregulates the activity of NKG2D ligands that are recognized by NK cells (21). As expected, PARP1 expression was higher in engrafting (CD34+) versus non-engrafting (CD34-) OCI-AML22 fractions (Fig 4A). Moreover, GSVA showed that genes known to be upregulated by NKG2D ligands were less enriched in CD34+ compared to CD34- fractions (Fig 4B and 4D). Inversely, genes downregulated by NKG2D ligands were more enriched in CD34+ compared to CD34- fractions (Figure 4C and 4D). Additionally, we have previously shown that LSCs exhibit a higher integrated stress response activity driven by ATF4 as compared to their downstream progeny (20). Through GSVA on each of the OCI-AML22 fractions, we confirmed that genes known to be upregulated by ATF4 were significantly enriched in the engrafting CD34+CD38- fraction, but less enriched in downstream progeny (Fig 4A and Supplementary Figure S7A), recapitulating our previous observations from primary AML samples. To directly track ATF4-driven induced stress response (ISR) activity across the differentiation trajectory from the CD34+CD38- fraction towards downstream progeny *in vivo*, we used a lentivector reporter to visualize ATF4 transcriptional activity (20) at single cell resolution by flow cytometry. The ATF4-reporter transgene ratio of each cell (GFP/BFP) reflects its relative ATF4 transcriptional activity. Isolated CD34+CD38- cells were transduced with this reporter and injected into mice (Figure 4B). Eight weeks after injection in mice, ATF4 activity was assessed in the CD34+CD38- and CD34+CD38+ populations generated in the xenografts (Fig 4B-C). The highest ATF4-driven ISR activity was detected in the CD34+CD38- fraction (Fig 4). This confirmed that the OCI-AML22 hierarchy recapitulates the ISR previously described in primary AML samples. Moreover, these results demonstrate that the OCI-AML22 LSC population can be readily modified with lentivectors. Notably, not all primary AML samples can be lentivector transduced, increasing the power of the OCI-AML22 model for functional investigations of stem cell driving features. To demonstrate the ability of OCI-AML22 to be valuable for future stemness studies, especially those focused on non-coding regulatory elements whose role in cell identity is now recognized (16,42), we undertook proof of concept that the OCI-AML22 LSC fraction can be CRISPR edited. We used a validated control olfactory gene (OR2W5) (43) to edit the OCI-AML22 CD34+CD38- fraction, then assessed their stemness potential in xenograft assays (Figure 4H). The CD34+CD38- fraction was efficiently edited (Figure 4J, (Supplementary Figure S8A) and retained the capacity for differentiation, cell expansion (Supplementary Figure S8B) and long-term repopulation capacity in NSG mice (Figure 4J-K).

**Figure 4:**
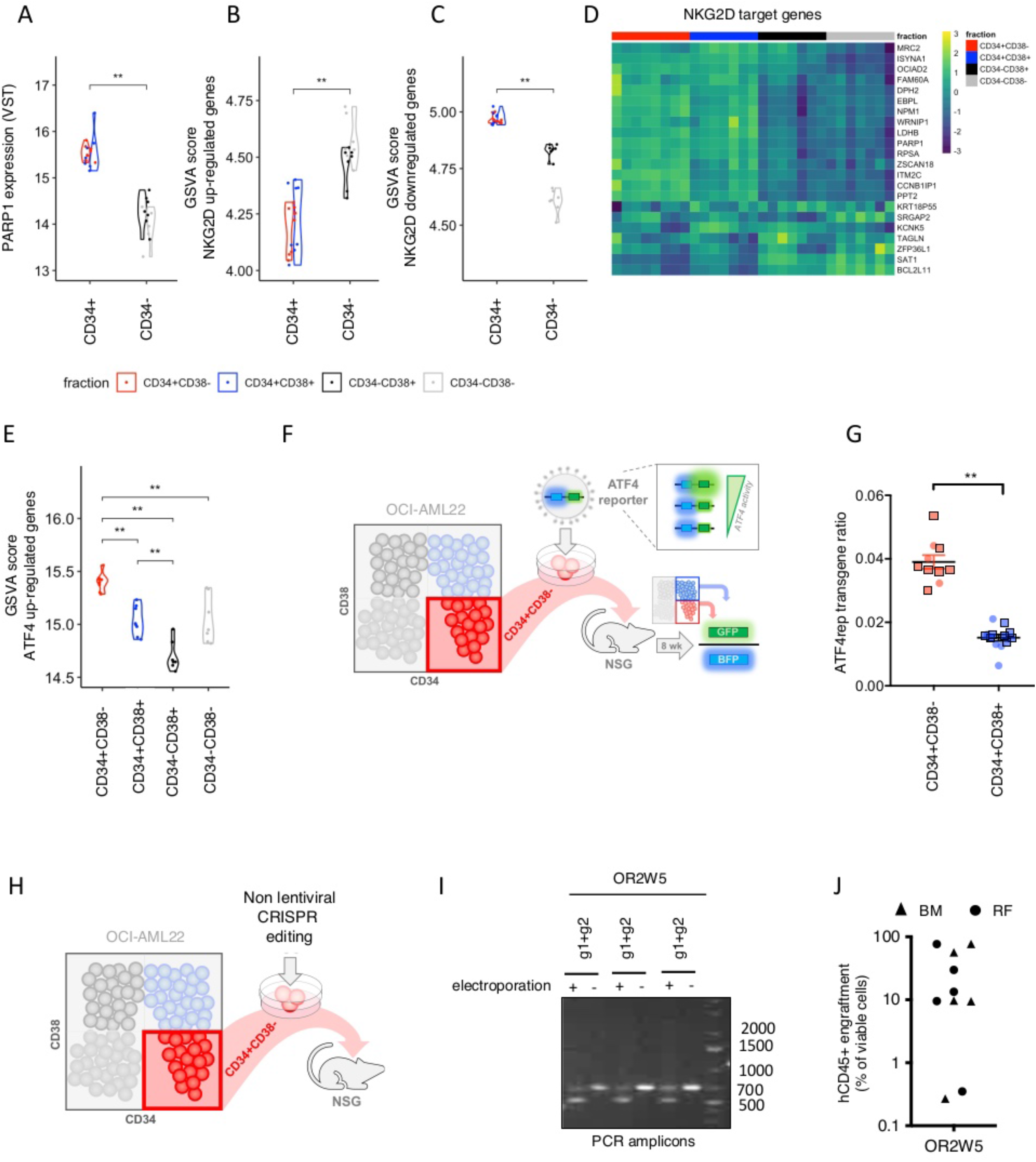
Efficient genetic modification of OCI-AML22 for functional interrogation of leukemia stemness. **A.** PARP1 expression of OCI-AML22 fractions was determined by RNA-Seq. **B-C**. GSVA score for NKG2D upregulated (B) or downregulated (C) genes as described in (21) across OCI-AML22 sorted fractions as indicated in Fig 2A. **D**. Supervised heatmap clustering of NKG2D target gene expressions of sorted OCI-AML22 fractions. **E.** GSVA score of ATF4 upregulated genes as described in (20) across individual OCI-AML22 fractions as extracted in Fig2B. **F**. Experimental scheme. **G.** OCI-AML22 CD34+CD38- cells were transduced with ATF4 lentiviral fluorescent reporter (ATF4rep) to track ATF4 transcriptional activity. BFP and GFP were assessed on viable human CD45+ cells extracted from the injected femora of NSG mice injected with 100k (round) or 200k (square) transduced cells, 8 weeks post xenotransplantation. The ATF4rep transgene ratio (GFP/BFP; BFP as internal transduction control) was then calculated for each indicated fractions of the 7AAD-AnnexinV- hCD45+ population from xenografts. **H.** The OCI-AML22 LSC fraction was CRISPR edited then injected in NSG mice to assess engraftment 21 weeks after injection. **I.** Gel electrophoresis of the amplified PCR fragment that corresponds to the OR2W5 edited locus flaked by g1 and g2 RNA guides, in the OCI-AML22 CD34+CD38- fraction. It allows visualization of the deletion of a 200 bp fragment within exon 1 of OR2W5 (uncut amplified PCR fragment:700bp, cut amplified fragment: 500 bp). **J**. CRISPR edited OCI-AML22 CD34+CD38- cells were injected in NSG mice. Representatif FACS profile (J) and engraftment level (K) was determined 21 weeks after.

Collectively, these data validate that OCI-AML22 can be used as a tool to study LSCs and interrogate stemness properties as previously reported for primary AML samples, both *in vitro* and *in vivo*. Moreover, the ability to genetically modify the LSC fraction of OCI-AML22 will broaden its usage for genetic screening and functional studies.

### The OCI-AML22 LSC-enriched cell fraction is composed of distinctly fated cells

To determine if the pool of LSC in OCI-AML22 was homogeneous or if it is composed of LSC subsets that varied in functional properties such as proliferation, differentiation and clonal maintenance, we analyzed individual cell fates within the LSC-enriched population. We adapted a single cell functional assay previously used for normal hematopoietic cells to study AML cells at the single cell level (44). Briefly, OCI-AML22 CD34+CD38- single cells were deposited into 96 wells plates pre-seeded with MS5 stromal cells (see Material and methods section) (Fig 5A). Flow cytometric analysis was performed after 7-10 days of *in vitro* co-culture to assess colony initiating potential, proliferative output and capacity of individual cells to completely or partially regenerate the OCI-AML22 cellular hierarchy (Fig 5A). We monitored the proliferative output of cells able to generate a colony and the differentiation properties of the progeny of the originally seeded cell. This assay revealed that a large proportion of CD34+CD38- cells were able to generate a colony (Supplementary Figure S9A). Interestingly, this single cell assay also revealed that even within seeded cells that were able to maintain and expand the CD34+CD38- fraction, both the differentiation potential and number of cells generated by each seeded cell were highly heterogeneous and formed a gradient between poorly differentiated colonies to colonies that have already regenerated the entire phenotypic hierarchy of OCI-AML22 (Supplementary Figure S9B-C). We classified initially plated cells as Alpha-type when they generated colonies that retained a high (>80%) percentage of primitive CD34+ cells, while Beta-type were defined as cells generating colonies already displaying the entire OCI-AML22 cellular hierarchy (Fig 5B). Alpha-type cells generated colonies exhibiting the highest percentage of CD34+CD38- (Supplementary Figure S9D) whereas Beta-type cells generated colonies with the highest percentage of more differentiated CD34-CD38+ and CD34-CD38- downstream cells (Supplementary Figure S9F-G). Beta-type cells represented only a minor proportion of the OCI-AML22 LSC fraction (Fig 5C and Supplementary Figure S9B). Nonetheless, Beta-type cells generated the largest colonies as measured by total cellular output per colony (Fig 5D) mainly due to a higher production of downstream CD34+CD38+, CD34-CD38+, and CD34-CD38- fractions (Fig 5E-H).

**Figure 5:**
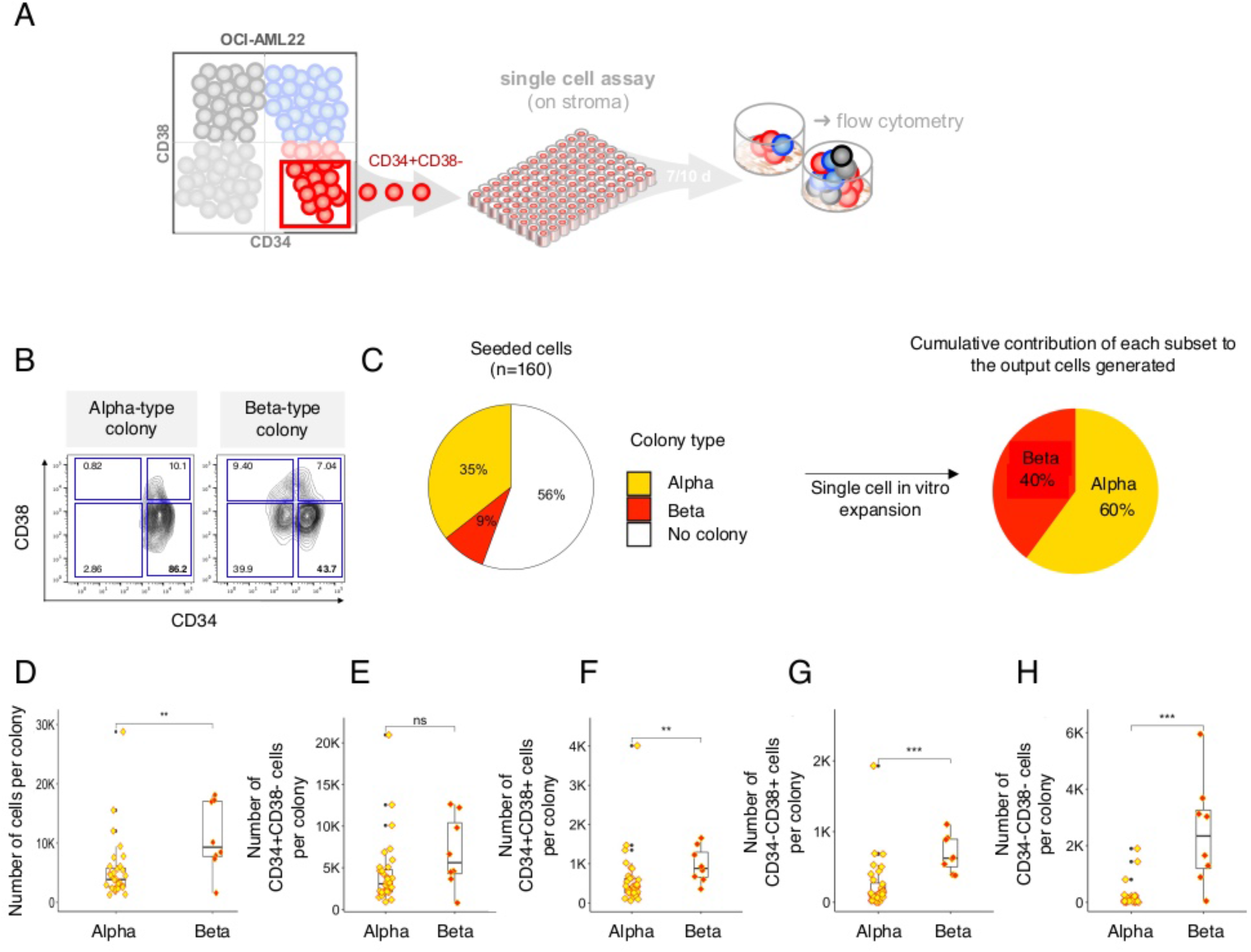
The OCI-AML22 LSC fraction is composed of distinctly fated cells. **A**, Experimental scheme. **B**, Extreme immunophenotypic profiles of Alpha-type and Beta-type colonies identified from single cell in vitro assay. **C**, Pie chart indicating the percentage of CD34+CD38- OCI-AML22 cells that have the capacity to generate either Alpha-type colonies, Beta-type colonies or do not generate any colony (left). Pie chart representing the cumulative contribution of Alpha or Beta type colonies to the overall cellular output generated (right). **D-H**, Absolute number of total cells (D), CD34+CD38- cells (E), CD34+CD38+ cells (F), CD34-CD38+ cells (G) or CD34-CD38- cells (H) per colony type generated from each single CD34+CD38- OCI-AML22 cells co-cultured with MS5.

Overall, this single cell *in vitro* assay highlighted that the CD34+CD38- OCI-AML22 fraction was heterogeneous and formed a gradient composed of at least 2 types of clonogenic cells. Both types were able to expand the LSC population but differed in their differentiation and proliferative potential. Interestingly, despite contributing to less than 10% of the OCI-AML22 CD34+CD38- compartment, Beta-type cells were responsible for 40% of the overall cellular output (Fig 5C) due to their greater ability to generate progeny (Fig 5F-H). This highlights the importance of studying AML cells at the single cell level since each cell contributes in a different way to the bulk output.

### Single cell ATAC-Seq identifies epigenetically distinct LSC types

To further explore the OCI-AML22 LSC fraction for distinct epigenetic stem cell states that would reflect distinct types of LSCs, we sorted the LSC fraction and performed single cell ATAC-Seq (Figure 6A). Using 8,911 cells which passed QC across 3 replicates, Uniform manifold approximation and projection (UMAP) was used to project per-cell Z score enrichments for accessibility of binding sites for all transcription factors in the ReMap 2020 project (45) calculated using chromVAR (Figure 6A, left UMAP). Our prior studies showed that throughout normal hematopoietic differentiation cells transition from being highly enriched for a LT/HSPC signature, indicative of dormant and high self-renewing stem cells, to stem cells with higher cellular output capacity, lower levels of quiescence and lower self-renewal capacity that exhibit a progressive depletion of the LT/HSPC signature coincident with progressive enrichment for an Act/HSPC signature (42). We investigated the enrichment of the entire catalogue of 10 chromatin accessibility signatures spanning sites predictive of different stages of normal hematopoiesis (42). LT/HSPC enriched single cells were distributed along a continuum of single-cells (Figure 6A, left UMAP). To further enrich those OCI-AML22 cells likely to have the highest self-renewal potential and hence representing putative LSCs, we selected 4,635 cells with LT-HSPC signatures (Z-score>0) and repeated the previous analysis upon this subset. (Figure 6A, right UMAP). We then used a Gaussian mixture model to identify 9 clusters (Figure 6B). To extract biological features distinguishing these clusters, we compared the Z-scores for chromatin accessibility signatures spanning the normal hematopoietic system among the cells of each cluster. (Figure 6C-D and Supplementary Figure S10A). These Z-Scores gradually changed between the clusters, suggesting the existence of distinct types of LSCs along a continuum. (Figure 6C-D and Supplementary Figure S10A). We then used slingshot to infer developmental trajectories over these clusters. This analysis revealed the existence of an axis with 2 extremes: from the green/red clusters on one end to the orange/blue clusters at the other end (Figure 6B, E-F and Supplementary Figure S10B). We split the longest trajectory into thirds to test for statistically significant differences in the enrichment of hematopoietic signatures (42) between different stages of the inferred trajectory (Figure 6F and Supplementary Figure S11A). This showed a significant progressive enrichment for the Act/HSPC signature from the first part of the trajectory, with significant progressive depletion for the mature, terminally differentiated signatures towards the end (Figure 6E and Supplementary Figure S10A-B and S11A). Overall, the portion of the OCI-AML22 LSC fraction most enriched for the LT/HSPC signature, is made of cells that form a gradient with 2 extremes. On one end, LSCs associated with a weaker Act/HSPC signature and potentially linked to lower progeny generation, while the other end contained LSCs associated with a higher Act/HSPC signature and potentially linked to higher progeny generation. This chromatin accessibility data is highly reminiscent of the functional differences identified from the in vitro single cell assays shown in Figure 5. Together, these two independent approaches suggest the existence of epigenetically distinct types of LSC within the CD34+CD38- OCI-AML22 fraction.

**Figure 6:**
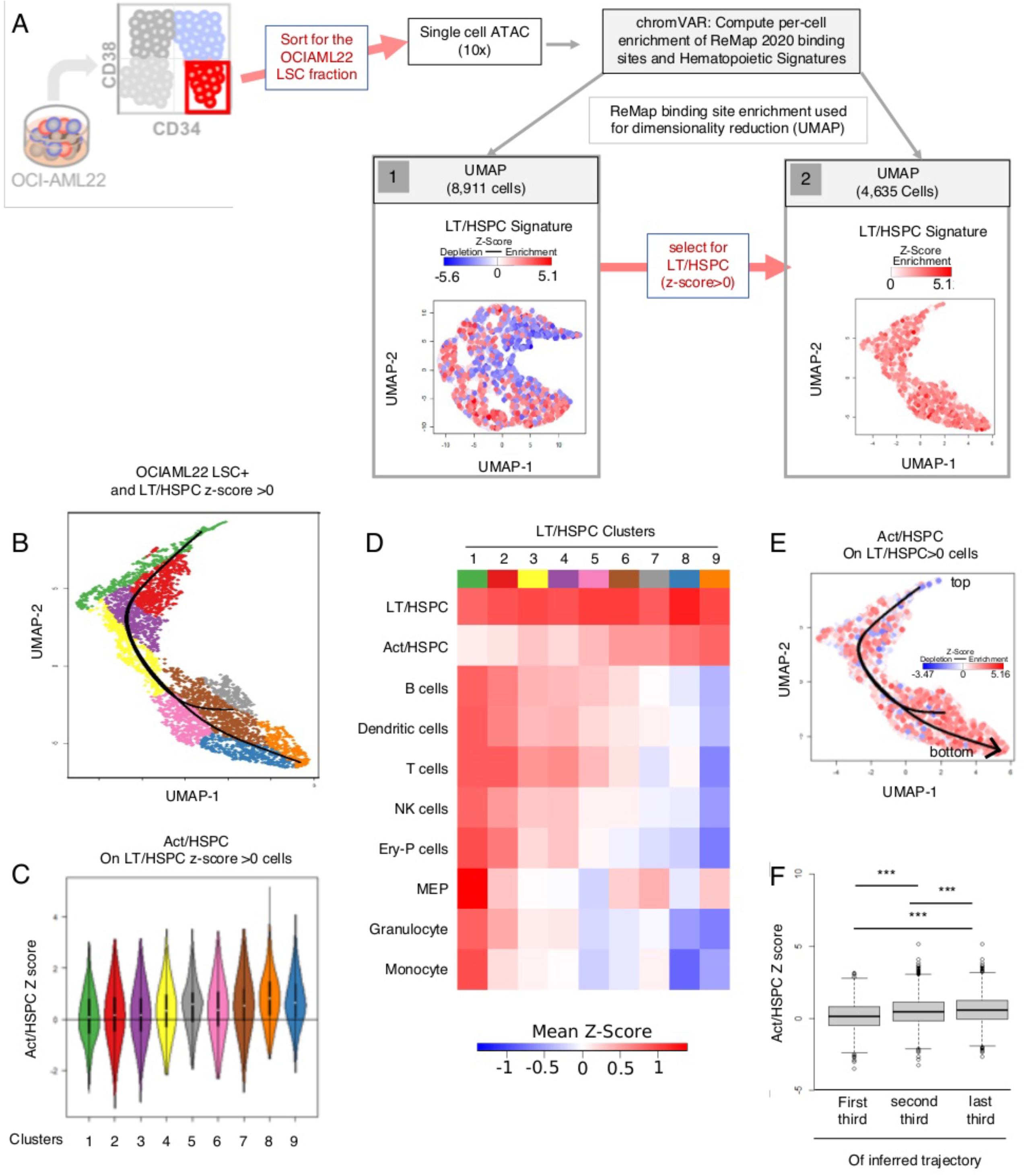
Single cell ATAC-Seq identifies epigenetically distinct LSC types. **A**, Schematic of experimental and computational pipeline. OCI-AML22 bulk cells cultured for about 3 months, were sorted for the LSC+fraction (CD34+CD38- cells) to perform single cell ATAC-Seq. Enrichment score for each of 10 chromatin accessibility signatures spanning normal hematopoiesis (42), and all binding sites from factors in the ReMap 2020 Project (45) was calculated for each single cell. Harmony was applied to the first 20 components of a principal components analysis to remove batch corrections, and UMAP was used to generate a 2D mapping of the batch corrected data (left). Cells with a Z-score>0 for the LT/HSPC signature were subsequently selected and UMAP was performed as previously described, on this subset (right). In each plot the colour indicates the per-cell Z-Score for the LT/HSPC signature, where blue indicates depletion, red enrichment. **B**, A gaussian mixture model was used to identify 9 clusters of cells from the LT/HSPC >0 UMAP representation (5A, left UMAP) and slingshot was used to infer a trajectory through the clusters. Cells in this plot are coloured according to cluster, while the inferred trajectory generated by slingshot is represented as a black line. **C**, Z-score enrichment for the Act/HSPC signature (42) calculated for the selected OCI-AML22 cells as described in 5A, left UMAP. **D**, Heatmap clustering of the mean Z-score for the enrichment of Takayama et al signature (42) that correspond to each signature within each cluster is represented. E, The LT/HSPC >0 UMAP representation (5A) colored based on enrichment for the Act/HSPC signature. The inferred trajectory generated by slingshot is represented as a black line. F, Single cells were split into three groups based around their position in pseudotime along the longest inferred slingshot trajectory. boxplots of Z-scores of enrichment for the Act/HSPC signature (top third: 1453 cells, middle third: 1453 cells, last third: 1452 cells. p values were calculated using a Wilcoxon signed rank test with FDR correction).

### The OCI-AML22 LSC fraction displays both latent and rapidly repopulating LSCs

To gain independent evidence for the existence of distinct repopulating cells within the LSC OCI-AML22 fraction, we used longitudinal limiting dilution xenografting to capture distinct repopulation kinetics. Since our in vitro single cell assay showed the presence of a small population within the OCI-AML22 LSC fraction that is capable of generating large colonies, we hypothesized that if such cells possessed repopulating ability in xenografts, they should generate larger grafts at earlier time points. By contrast, cells that yielded smaller colonies in vitro would be predicted to have lower proliferative capacity in xenografts. Mice were injected with different cell doses to reach a cell dose averaging the limiting dilution dose (30k, 7500, 1875, 469 cells per mice) (Figure 7A). The ability of LSCs at limiting dose to generate detectable engraftment was determined at 2 time points: 8 weeks and 12 weeks after injection of the same batch of sorted OCI-AML22 CD34+CD38- cells (Figure 7A). Twelve weeks after injection with the lowest number of cells (469 cells per mice), we observed that engraftment level could be detected in 60% of the mice injected (Figure 7B, outer circle) confirming that the number of cells injected must be close to the limiting dilution dose. However, when the efficiency of engraftment of this low cell dose group was assessed at 8 weeks, only 33% of mice were engrafted (Figure 7B, innercircle). This result suggests that LSCs with engraftment potential must be present at 8 weeks, but remain below the detection level by flow cytometry, since such LSC eventually are able to generate a detectable graft at 12 weeks. These clonal studies suggest the existence of 2 types of LSCs with different repopulation kinetics. We observed similar results in mice injected with 1875 cells (Figure 7C). While engraftment could be detected at 12 weeks for all the mice injected at this cell dose, we again detected grafts early (at 8 weeks) for only a portion of the mice in this cohort (66.6%) (Figure 7C). In line with this result, the estimated LSC frequency at 8 weeks after injection was lower than at 12 weeks (1/1477 vs 1/286, respectively; p=0.012, (Figure 7D). At non-clonal cell doses we expect that all xenografts are repopulated from both slow and rapid repopulating LSCs (Supplementary Figure S12A). Collectively, these results establish that the CD34+CD38- OCI-AML22 fraction is functionally heterogeneous with distinct rapid and slowly repopulating LSCs.

**Figure 7:**
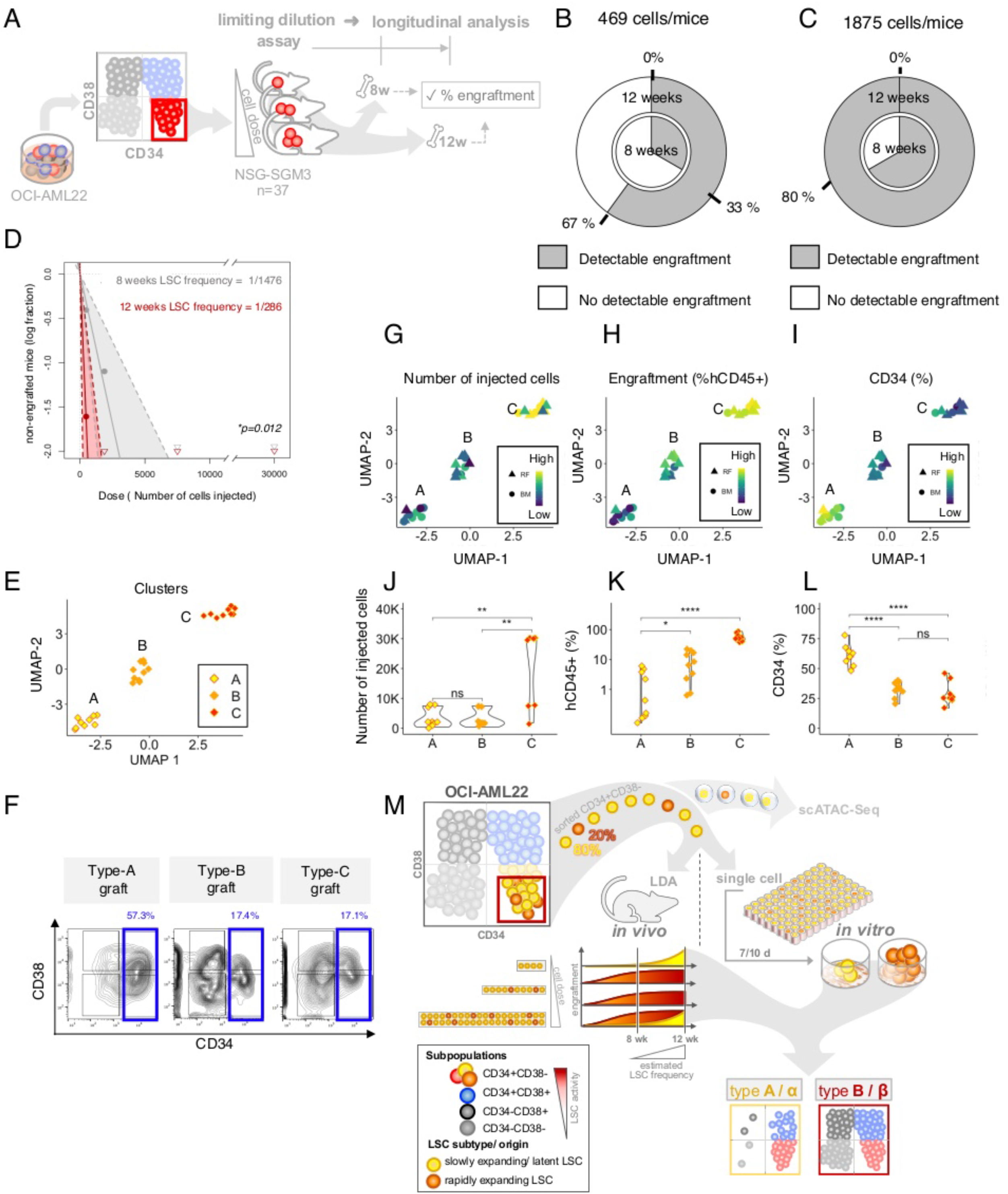
The OCI-AML22 LSC fraction displays latent and rapidly repopulating LSCs linked to distinctly fated outcomes. **A**. Schematic representation of the experiment. NSG-SGM3 mice were injected intrafemorally at limiting dilution with the indicated cell doses of CD34+CD38- OCI-AML22 cells. Engraftment and differentiation profile was assessed 8 and 12 weeks after injection. **B.** Pie chart representing the percentage of mice for which engraftment could be detected at 8 weeks (inner circle) or 12 weeks (outer circle), after injection of 469 OCI-AML22 CD34+CD38- cells sorted from a bulk culture expanded for 3-4 months in vitro. **C.** Pie chart representing the percentage of mice for which engraftment could be detected at 8 weeks (inner circle) or 12 weeks (outer circle), after injection of 1875 OCI-AML22 CD34+CD38- cells sorted cells. **D.** Calculated LSC frequency at 8 or 12 weeks after injection of CD34+CD38- OCI-AML22 cells at doses indicated in Supplementary Figure S10A. **E-I** UMAP-based clustering of engraftment parameters derived from non-injected bone marrow (BM) and injected femur (RF) (parameters: total human cell counts, CD34 expression, engraftment level, number of injected cells per mice) of NSG-M3S mice after injection of CD34+CD38- OCI-AML22 cells at cell doses per mouse indicated in Supplementary Figure S10A. Xenografts are colored based on (E) clusters determined using the k-means clustering method. (G) number of injected cells per mice, (H) human engraftment level (hCD45%), (I) CD34 percentage (of hCD45+). **J-L**. Statistical analysis for each UMAP cluster showing number of injected cells (J), engraftment level (K), percentage of CD34+ cells (L). **F**, Representative immunophenotypic profiles of type-A,B or C grafts based on CD34 and CD38 cell surface expression (hCD45+ subgated cells are shown).

### Individual LSCs within the OCI-AML22 CD34+CD38- fraction are linked to distinctly fated outcomes

To gain deeper understanding of the functional heterogeneity observed within the pool of OCI-AML22 LSCs, we carried an integrated analysis of all the functional data we collected on engraftment and differentiation parameters at the 12-week time point. We performed multi-dimensional data reduction using Uniform Manifold Approximation and Projection (UMAP) to integrate engraftment level and cell counts as a surrogate for repopulation capacity and the distribution in percentages across the 4 subpopulations as a surrogate for differentiation capacity (Figure 7E-I). The UMAP analysis revealed 3 clusters (Figure 7E, clusters A, B and C) that described distinct graft types (Figure 7F and Supplementary Figure S12B). Cluster C was composed of grafts predominantly generated from the highest cell dose injected (Figure 7G and 7J) that also resulted in the highest engraftment levels (Figure 7H and 7K). In contrast, cluster A and B were composed of grafts generated by similar number of injected cells (Figure 7G and 7J), but differed by the size of the graft generated (Figure 7H and 7K) and the differentiation potential of cells injected, as assessed by the remaining CD34 expression of their resulting progeny (Figure 7I and 7L). Additionally, cell doses injected in cluster A and B were low and approaching a limiting LSC frequency dose. Thus, the initial injected cells that generated grafts found in cluster A (type-A grafts) were characterized by the slowest repopulating capacity while still maintaining the most primitive phenotype. Conversely, the initial injected cells that generated grafts found in cluster B (type-B grafts) exhibited higher repopulation and differentiation capacity as compared to those generating type-A grafts. This observation suggested that the OCI-AML22 LSC fraction contains at least two functional types of LSCs differing in their repopulation kinetic and differentiation potential. These 2 types are both functional LSCs since each is able to generate a graft in immunodeficient mice and expand the LSC fraction *in vivo* (Supplementary Figure S12C), fulfilling the gold standard for defining LSCs (9,15). Altogether, our functional study shows that the OCI-AML22 LSC fraction is composed of at least two functional types of LSCs with distinct repopulation kinetics: primitive-like LSCs that slowly repopulate the bone marrow of immunodeficient mice, generating type-A grafts; and differentiation-prone, rapidly repopulating LSCs generating type-B grafts (Figure 7M).

## Discussion

We developed a primary AML derived stem cell model, OCI-AML22 to address difficulties in obtaining and extracting functional LSCs from primary AML samples and that represents the bottleneck for stemness related studies. OCI-AML22 recapitulates the hallmarks and cellular states of primary AML samples. In particular, the LSC fraction closely recapitulates, at the functional, transcriptional and epigenetic level, stem cell features shared by the largest panel of functionally assessed primary stem cell fractions available in the literature (8,9,21,42). These common stem cell properties are prognostic across multiple cohorts of AML patients, collectively spanning more than 1000 AML patients (9,17,46) demonstrating the clinical relevance of the OCI-AML22 model.

In this model, LSCs are easily identified, enriched and expanded while maintaining primary stem-like features. This simplicity, together with the ability to genetically modify the LSC fraction with lentivectors and CRISPR editing strategies contrasts with the difficulties the research community is facing with primary AML samples. Any assay requiring large numbers of stem cells (e.g. mass spectrometry based proteomics and metabolomics), as well as drug or genetic high throughput stemness-based screens are now feasible. Moreover, CRISPR editing strategies have opened new avenues beyond only gene interrogation including editing non-coding elements that are increasingly being recognized as being stemness and/or cancer regulators. However, while CRISPR editing has been achieved in bulk traditional AML cell lines and optimized in primary human HSCs (43,47), it has never been reported, to our knowledge, in either human LSC or even primary AML samples. OCI-AML22 shares some key features with OCI-AML8227, another AML model we previously developed, including organization into a phenotypic and functional hierarchy (EL, in preparation, (22). However, the LSC fraction is less enriched in OCI-AML8227. Both models closely recapitulate primary AML features (EL, in preparation), unlike traditional AML cell lines and provide multi-omic datasets that represent a rich and unique resource to study stemness in AML.

Compared to the other traditional AML cell lines, we showed here that the OCI-AML22 LSC fraction maintains both the genetic and functional heterogeneity characteristic of LSCs found in primary AML samples. First, through WGS, we showed that OCI-AML22 maintains at least two genetic subclones throughout *in vitro* and *in vivo* expansion that can be traced back to minor subclones found in the donor sample. Second, combining functional, phenotypic and epigenetic outputs on the OCI-AML22 LSC fraction, we further discovered functional heterogeneity within the LSC compartment showing the existence of latent and rapid repopulating stem cells. These results support prior studies of primary AML samples that suggested such LSC subclasses might exist (11,40,48) and are highly reminiscent of the latency phenomenon we reported for both LSC and normal HSC obtained from human primary samples. Despite being proposed almost a decade ago from lentivector-mediated clonal tracking of primary AML samples in longitudinal xenografts (38), it has been difficult to study the functional heterogeneity of the LSC compartment of primary samples due to LSC scarcity (15). The OCI-AML22 model overcomes this limitation. Our WGS study showed that these 2 functional types of LSCs are >99% genetically similar, only differing in a small deletion of chromosome 11. This argues for the concept that these two LSC classes arise from epigenetic processes. However, our genetic analysis is not deep enough to rule out the possibility that some genetic differences between these subsets, including the small 11q deletion, are functionally important. Our identification of epigenetically distinct LSC subsets is of particular interest since epigenetic processes have been shown to be both clinically relevant in cancer (3,4,33,49–55) and reversible (56). This discovery, combined with the technical ability to deconvolute the OCI-AML22 LSC population at the genetic and functional level, pinpoints to the utility of OCI-AML22 as a unique model to resolve the considerable challenges linked to studying functional heterogeneity of the LSC compartment.

Ultimately, the availability of the OCI-AML22 model for functional validation, as well as the extensive datasets generated on different functionally characterized LSC and non-LSC fractions, will open up new avenues to investigate stemness processes and identify vulnerabilities. The ability to study LSCs at the functional level as well as to specifically deconvolute this compartment is essential for any future therapeutic target development. Indeed, it is likely that stem cells that are associated with different levels of differentiation and repopulation capacity may also have different responses to current therapies, potentially explaining why some therapies fail. Because of the difficulties to obtain both highly enriched populations of LSCs, and to genetically manipulate them, it has been challenging to study and genetically manipulate them. OCI-AML22 serves to close this gap, while also working with a model that is clinically relevant.

## Acknowledgements

J.E.D is supported by funds from the: Princess Margaret Cancer Centre Foundation, Ontario Institute for Cancer Research through funding provided by the Government of Ontario, Canadian Institutes for Health Research (RN380110 - 409786), Canadian Cancer Society (grant #703212 (end date 2019), #706662 (end date 2025)), Terry Fox New Frontiers Program Project Grant (Project# 1106), a Canada Research Chair, Princess Margaret Cancer Centre, The Princess Margaret Cancer Foundation, and Ontario Ministry of Health. HB was supported by the Helena Lam Fellowship in Cancer Research Fund. We thank Olga Gan for technical support, Andy G X Zeng for helpful insight, and the Leukemia Tissue Bank at Princess Margaret Cancer Centre/ University Health Network as the source of the primary sample blood. We thank Nicholas Khuu from the Princess Margaret Genomics Centre, Toronto, Canada (www.pmgenomics.ca) for ATAC-Seq sequencing, Sergio Pereira of The Centre for Applied Genomics, The Hospital for Sick Children, Toronto, Canada for assistance with RNA sequencing. We thank Sabrina Smith and Gaayatiri Pushparaj for assistance with experiments. We thank Melissa Anders for providing support for writing, Eric Lechman and Jean C Y Wang for critical reading of the manuscript and all members of the Dick lab for critical feedback. This work is dedicated to the memory of A-M W.

## Author contributions

H.B. established the model, conceived the study, performed research, performed bioinformatic analysis and wrote the manuscript. A.Mu. and M.C-S-Y. performed bioinformatic analysis, SA.MT and M.C-S-Y. assisted with bioinformatic analysis. E. W. helped for methods, N.M. and A.Mi, performed research, M.L., provided funding, M.M. and A.A. provided AML samples, E. O. and F. N. supervised part of the study, A.Mu. and M.M. edited the manuscript, C.A. performed research, K.B.K. performed research, wrote the manuscript, provided input to conceive the study. J.E.D. wrote the manuscript, secured funding and supervised the study.

## Material and Methods

### Primary samples

Primary AML samples were collected with written informed consent according to the procedures approved by the University Health Network (UHN) Research Ethics Board (REB 01-0573-C). 400 000 cells were initially plated at 2 millions cells per ml for each tested sample. Cells were maintained in culture for the indicated time in the following medium: X-VIVO 10 (Lonza, BE04-380Q) supplemented with 20% BIT 9500 Serum Substitute (StemCell technologies, 09500), SCF (50 ng/ml) (Miltenyi Biotech, 130-096-696), IL3 (25 ng/ml)( Miltenyi Biotech, 130-095-069), TPO (50 ng/ml; Peprotech, 300-18), FLT3L (50 ng/ml; Peprotech, 300-19 B), IL6 (25 ng/ml; Miltenyi Biotech, 130-093-934), G-CSF (25 ng/ml; Miltenyi Biotech, 130-093-861).

### Clinical characteristics of the patient from which OCI-AML22 originated

Male, 58 old, diagnosed with acute myeloid leukemia with myelodysplasia related changes of unclassified FAB carrying TP53 mutation (variant C743G>A), VAF 0.939. Karyotype of the patient at diagnosis was 46,XY,del(3)(p21),+del(3)(q21),+del(6)(q21),del(7)(q22q32),-11,-14,-17,+mar[9].

#### Disease history of the donor that generated OCI-AML22 and OCI-AML22 establishment

The donor that generated the OCI-AML22 model had a primary refractory response to the classical 7+3 based induction chemotherapy combining daunorubicin and cytarabine. It was followed by a FLAG-IDA re-induction. After an initial clinical remission, the patient relapsed 55 days later. He then received NOVE HiDAC re-induction therapy, followed by haploidentical hematopoietic stem cell transplantation, which led to a second remission. He then relapsed 78 days after transplantation. Following this second relapse, the patient started a leukoreduction protocol based on azacytidine and hydroxyurea. At that point, a peripheral blood sample was collected and blasts were isolated to generate the OCI-AML22 sample. Later, the patient was treated with Myelotarg, but complete remission was not achieved. Overall survival was 480 days.

The peripheral blood from the donor was subjected to ficoll separation followed by red blood cell lysis. At that time, the percentage of blasts was 89%. Freshly obtained cells were then expanded in culture in the following medium, with final concentrations as indicated: X-VIVO 10 (Lonza, BE04-380Q) supplemented with 20% BIT 9500 Serum Substitute (StemCell technologies, 09500), 1x Glutamax Supplement (Thermo Fisher Scientific, 35050061), Primocin 0.1 mg/ml (invivogen, ant-pm-1), SCF (200 ng/ml;Miltenyi Biotech, 130-096-696), IL3 (20 ng/ml;Miltenyi Biotech, 130-095-069), TPO (20 ng/ml; Peprotech, 300-18), FLT3L (40 ng/ml; Peprotech, 300-19 B), IL6 (10 ng/ml; Miltenyi Biotech, 130-093-934), G-CSF (10 ng/ml; Miltenyi Biotech, 130-093-861). Cells were maintained at a density of 0.8×10^6^ cells/ml and passaged twice a week, in a 96 well flat bottom plate surrounded by PBS on the outside wells. A full medium change was done once a week. The CD34+CD38- fraction or CD34+ fraction was regularly sorted to serially expand the cells.

#### Cytogenetic of the donor that generated the OCI-AML22 model

The donor cells display a complexe cytogenetic that is characterized by:

- A series of deletions on chromosomes 4, 7, 8, 11.
- A series of amplifications on chromosome 11.
- Copy number alterations on chromosome 17. Of note deletion of chromosome 17 is also consistent with the TP53 mutation LOH observed on the diagnosis sample.
- A series of amplifications of chromosome 21.
- Partial chromosome Y deletion

#### Xenotranplantation

All animals used for this study were treated in accordance with institutional guidelines. Female and male NSG mice (NOD.Cg PrkdcscidIl2rgtm1Wjl/SzJ; The Jackson Laboratory) or male NSG-SGM3 mice as indicated in figure legends were irradiated with 250 rads the day before intrafemoral injection. Bulk cells expanded in vitro for an average of 4 months were used throughout this study unless specified otherwise and are considered the OCI-AML22 model. Fractions were obtained from expanded cells in vitro after an average of 4 months (unless specified otherwise), then sorted according to their CD34 and CD38 expression, using AnnexinV-FITC (556419, BD) and 7-AAD (559619, BD) to exclude apoptotic and dead cells. The number of injected cells per experiment is indicated in the figure legends. For lentiviral transduction experiment, cells were transduced with the ATF4 reporter as described in (57) in their standard culture medium overnight at a density of 0.8×10^6^ cells/ml, washed and injected the day after. Mice were euthanized and the injected femur (right femur /RF) and the non-injected left femur as a surrogate for engraftment in the bone marrow (BM) were flushed separately in MEMalpha with 10% FBS. Engraftment level was assessed by flow cytometry on a FACSCelesta instrument (BD) with the following antibodies: mCD45-FITC (553080, BD) or hCD45-FITC (561865, BD, clone HI30) combined with hCD45-APC-Cy7 (624072, BD, clone 2D1), hCD34-PE (348057, BD), hCD38-PE/Dazzle (303538, BD) and 7-AAD (559619, BD). For secondary engraftment, human cells collected from NSG engrafted mice were sorted using 7-AAD (559619, BD), AnnexinV-FITC (556419, BD), hCD45-APC-Cy7 (624072, BD), then injected into NSG-SGM3 mice at the indicated cell dose per mice and sacrificed at 8 weeks after. The engraftment level was determined as for the primary engraftment assay.

#### RNA-seq

Primary cells were expanded generating the OCI-AML22 model and collected at different time points to sort the resulting subfractions based on CD34 and CD38 cell surface expression as indicated in figure legends. Total RNA was extracted using the mirVana™miRNA Isolation Kit, with phenol (Invitrogen) as recommended. Samples that passed quality control according to integrity (RIN>8) and concentration as verified on a Bioanalyzer pico chip (Agilent Technologies) were subjected to further processing by the Center for Applied Genomics, Sick Kids Hospital. The SMART-Seq v4 Kit (SSv4) (Takera) followed by Illumina Nextera XT library prep was used as recommended by the manufacturer. Briefly, the same amount of RNA was used for library preparation. After cDNA conversion, cDNA was run on a bioanalyzer. 1ng of cDNA was used for the Nextera library preparation, then subjected to QC on a bioanalyzer and qPCR for the final library concentration. For sequencing, all libraries were pooled at equimolar amounts, followed the illumina protocol to dilute the pool to the appropriate concentration for sequencing. All samples were sequenced in parallel to avoid batch effects, on the NovaSeq 6000, S1 flow cell, PE100bp. Approximately 43 million paired reads per sample were generated.

#### RNA-seq analysis

Sequencing was aligned using STAR 2.5.2b (58) against GRCh38 and transcript sequences downloaded from Ensembl build 90. Default parameters were used except for the following: “--chimSegmentMin 12 --chimJunctionOverhangMin 12 --alignSJDBoverhangMin 10 --alignMatesGapMax 100000 --alignIntronMax 100000 --chimSegmentReadGapMax parameter 3 --alignSJstitchMismatchNmax 5 −1 5 5”. Counts were obtained using HTSeq v0.7.2. Variance stabilized normalized counts were generated. The variance stabilized normalized counts was generated by DESeq2 v1.22.2. Gene set enrichment was performed using GSEA PreRanked v3.0 (http://www.broad.mit.edu/gsea/) against the indicated gene sets as specified in figure legend (settings: method “ssgea, kcdf: “Gaussian”). The UMAP package in R (59) was used (parameters: number of neighbors = 5, min dist = 0.01) to reduce the dimensionality of (Figure 7G-I). Clusters were identified using the k means method. PCA has been generated using the plotPCA function from DESeq2 v1.22.2 package and the top 1000 variable genes. Graphic has been made using ggplot2.

#### Generation of references stem cell signatures used in Figure 4s

The normal karyotype LSC+ signature has been generated by taking the average gene expression of each of the 104 genes part of the LSC104 signature, using the 60 LSC+ fractions obtained from patients with normal karyotype and obtained from (9). The abnormal karyotype LSC+ signature has been generated by taking the average gene expression of each of the 104 genes part of the LSC104 signature, using the 55 LSC+ fractions obtained from patients that did not carry a normal karyotype and obtained from (9). The LSC frequency High signature was generated by taking the average gene expression of each of the 104 genes part of the LSC104 signature, using the 32 LSC+ fractions that presented the highest LSC frequency within our entire cohort of 166 functionally assessed fractions, obtained from previous work (9). The LSC frequency Med signature was generated by taking the average gene expression of each of the 104 genes part of the LSC104 signature, using the 52 LSC+ fractions that presented a medium LSC frequency within our entire cohort of 166 functionally assessed fractions, obtained from previous work (9). The LSC frequency Low signature was generated by taking the average gene expression of each of the 104 genes part of the LSC104 signature, using the 28 LSC+ fractions that presented a low LSC frequency within our entire cohort of 166 functionally assessed fractions, obtained from previous work (9). LSC frequencies for each of these categories are detailed in Supplementary Figure S5B, and are representative of a large panel of AML samples as presented in Supplementary Figure S6.

#### ATAC-Seq library preparation

Library preparation for ATAC-Seq was performed by the the “Princess Margaret Genomics Centre, Toronto, Canada (www.pmgenomics.ca) on 100,000 sorted CD34+CD38- OCI-AML22 cells that have been cultured for about 4 months, with Nextera DNA Sample Preparation kit (FC-121-1030, Illumina), according to previously reported protocol (60). Libraries for ATAC were sequenced on the Novaseq 6000 System (Illumina) to generate paired-end 50-bp reads. Reads were mapped against the hg19 human reference genome using BWA (61) with default parameters. All duplicate reads, and reads mapped to mitochondria, chrY, an ENCODE blacklisted region or an unspecified contig were removed (62). MACS 2.0.10 (63) was used to call peaks from aligned reads.

#### scATAC-Seq and Analysis

Cells from three sorted CD34+CD38- OCI-AML22 replicates were processed on the 10X Genomics single cell ATAC-seq platform, and 8,911 single cells were retained based on default cellranger QC criterion and chromVAR depth filtering (64). chromVAR (with default settings) (64) was used to calculate the enrichment of each of the hematopoietic signatures identified in (42) in each single cell, as well as for the catalogue of ChIP-Seq sites of specific transcription factors identified in the ReMap project (45). Harmony (65) was used to batch correct the first 20 components of a principal components analysis (PCA) performed on the Per-cell enrichment of all human TFs identified in the ReMap project (45), and the UMAP package in R (59) (with default parameters except number of neighbours equal to 14) was subsequently used to produce a 2D mapping from the batch corrected PCA. This step was repeated for all single cells with LT/HSPC signature enrichment >0 (with number of neighbours in UMAP set to 7). The Mclust R package with default parameters was used to identify clusters from the UMAP dimensionality reduction via gaussian mixture modelling. Slingshot was used with default parameters to infer possible developmental trajectories through these clusters.

#### In vitro single cell assay

MS-5 stromal cells were kindly gifted from Dr. K. Itoh from Japan (66). MS-5 cells were seeded into 0.2% gelatin coated 96 well-flat bottom plates (Thermo Fisher, 167008) at a density of 4-5×105 cells per well in Myelocult medium (H5100, Stem Cell Technology). In the morning, the medium was changed using OCI-AML22 culturing medium described before (50uL/well). The CD34+CD38- OCI-AML22 fraction from cultured cells (about 3-4 months) was sorted and directly plated in 96 well plates precoated with MS5, then refilled with 50uL of fresh medium per well. 7 to 10 days after, half of each well was collected, washed and stained with hCD45-FITC (561865, BD), hCD34-APC-Cy7 (624072, BD), hCD38-APC (340439, BD). 7-AAD (559619, BD) or sytox blue (S11348, thermoFisher) were used to exclude dead cells, then acquired on a FACSCelesta (BD). Wells for which more than 10 viable cells could be detected in the hCD45+ viable cells gate were identified as a colony. To determine the percentage of CD34/CD38 within the colony, a minimum number of 300 cells in the hCD45+ viable cells gate was set to increase confidence in the percentage.

#### Lentiviral production of ATF4 reporter

The ATF4 reporter was used as previously reported (20). Pseudotyped lentiviral particles were produced and titers calculated as previously described (57). Lentivirus were concentrated 100x by ultracentrifugation, resuspended in X-VIVO 10 supplemented with 1% BSA and stored at – 80C until use. Cells were transduced overnight and washed the day after with a fresh medium before their injection in mice.

#### CRISPR editing of the OCI-AML22 LSC fraction

The protocol was adapted from (43) for CRISPR editing of LSCs. In PCR sterile tubes, crRNA synthetized from IDT as Alt-R CRISPR/Cas9 crRNA and targeting the control OR2W5 olfactory gene using 2 guides: crRNA-g1_OR2W5 (GACAACCAGGAGGACGCACT) and crRNA-g2_OR2W5 (CTCCCGGTGTGGACGTCGCA) where annealed with tracrRNA(IDT, 1072533) in a ratio 1crRNA g1, 1crRNAg2, 2 tracrRNA, at 95 °C for 5 min, then cooled down to room temperature. Then 1.2uL of the resulting functional gRNA duplex was combined with 1.7uL cas9 protein (IDT, 1081059) and 2.1uL PBS and incubated for 15 min at room temperature. 1uL of electroporation enhancer (IDT, 1075916) was subsequently added in each reaction to generate the complex mix. In parallel, cultured OCI-AML22 cells, pre-sorted the day before for its CD34+CD38- fraction using the MoFlo XDP U/VBR Beckman Coulter sorter, and incubated overnight in their own medium, were washed with room temperature PBS using a benchtop centrifuge at 3rpm for 2 minutes. 50 000 to 100 000 cells per reaction was then resuspended in 20uL of P3 solution (Lonza, V4XP-3032) and mixed by pipetting with the full gRNA-Cas9 complex mix previously assembled then added in the electroporation chamber well (Lonza, V4XP3032). Cells were electroporated with the program EO-100 using the Lonza Nucleofector. Then, 180 μl of pre-warmed OCI-AML22 media (as described above) was added. Cells were transferred in a 96 plate well surrounded by PBS to maintain humidity, then left overnight in the incubator before replacing 100uL of old medium with fresh OCI-AML22 media. After each CRISPR/Cas9 RNP electroporation, a small subset of cells was cultured in the OCI-AML22 media for about a week to obtain enough cells for DNA analysis. Genomic DNA was isolated from bulk cells using the Agencourt GenFind V2 (Beckman Coulter, A41499). The CRISPR/Cas9 engineered genomic locus was amplified via PCR as previously described (43). For each PCR reaction, 23 μl of eluted genomic DNA was mixed with 1 μl of forward and reverse primer (10 μM) and 25 μl of AmpliTaq Gold 360 Master Mix (ThermoFisher, 4398881). The PCR program was: 95 °C for 10 min, followed by 95 °C for 30 s, 56 °C for 30 s and 72 °C for 1 min (40 cycles) and then 72 °C for 7 min.PCR primers: Control (OR2W5) forward primer: 5’-TCGGCCTGGACTGGAGAAAA-3’, Control (OR2W5) reverse primer: 5’-GAGACCACTGTGAGGTGAGA-3’. A portion of the PCR amplicon was run on an agarose gel to verify both PCR specificity and efficiency at deleting a 200 bp fragment within exon 1 of OR2W5 (uncut amplified PRC fragment:700, cut amplified fragment: 500 bp) (see gel, Figure 4I). The rest of PCR products were purified using the MinElute PCR Purification Kit (Qiagen, 28004). Sanger sequencing was performed using the OR2W5 forward PCR primer (5’-TCGGCCTGGACTGGAGAAAA-3’). Chromatograms were analyzed using the online tool TIDE (https://tide.deskgen.com/)(67) to verify that the break site is at the expected location which validates precision of the cut (Supplementary Figure S7A).

#### Whole genome sequencing analysis

We used 2 independent pipelines: HMMcopy (https://bioconductor.org/packages/release/bioc/html/HMMcopy.html) and celluloid (68), to detect copy number abnormalities. The whole genome sequencing data was aligned to hg38 using bwa v0.7.12 with default settings. Reads were sorted and duplicates were marked with picard v2.21.4. Copy number calls were made using both HMMcopy v0.1.1 and celluloid v0.11.7 with 1kb windows. Telomeric and centromeric regions were removed from further analysis. We considered anything with a copy number below 1.5 as a loss and greater than 2.5 as a gain. Using bedtools v2.29.2 we identified overlapping regions in pairwise fashion to determine the percentage of overlap between each sample. To determine if subclones were present, we focused on the alterations found on region of chr11 q arm. Using the same segments identified in bulk_1m, we compared the distribution of the binned copy number values between adjacent segments using a wilcox test. This was done in all samples to identify if there was a small change between segments that were not identified by the copy number callers.

#### Statistical analysis

GraphPad Prism or R was used. Unless specified, Mann Whitney test was performed. *<0.05, **p<0.01, ***p<0.001.

## Tables legends

**Table 1**: Clinical characteristics of AML patient samples tested for in vitro expansion from Supplementary Figure S1A.

**Table 2**: Limiting dilution assay at 12 weeks. OCI-AML22 cells were expanded in culture for 4 months before CD34+CD38- cells were isolated and injected into NSG-SGM3 mice at the indicated cell doses. The number of injected mice (tested) and engrafted (response) as well as lower, estimated and upper LSC frequencies are indicated 12 weeks after injection.

**Table 3**: Chromosome 11q fragments being compared coordinates. Segment regions that were detected using hmmcopy in the cultured sample and were used to break down the chr11 arm q of the donor sample or the fractions sorted from xenografts.

**Table 4**: Comparisons of fragments between the donor or xenografts fractions ( CD34+ or CD34-) and the cultured sample. Segment regions that were detected using hmmcopy (Table 3) in the cultured sample were used to break down the chr11 arm q of the donor sample, or xenografts into matching regions. Within these segments, each dot (at 1kb intervals) was taken to compare the average copy number to the adjacent segments and thus determine if the average copy numbers were different between the indicated adjacent regions. wilcox tests were run. significance shows the existence of amplifications that can be detected at a subclonal level in the interrogated sample: donor sample or xenografts (CD34+ fraction, CD34- fraction or both)

## References

1. Papaemmanuil E, Gerstung M, Bullinger L, Gaidzik VI, Paschka P, Roberts ND, et al. Genomic Classification and Prognosis in Acute Myeloid Leukemia. N Engl J Med. 2016;374:2209–21.

2. Brewin J, Horne G, Chevassut T. Genomic landscapes and clonality of de novo AML. N Engl J Med. 2013;369:1472–3.

3. Cancer Genome Atlas Research Network, Ley TJ, Miller C, Ding L, Raphael BJ, Mungall AJ, et al. Genomic and epigenomic landscapes of adult de novo acute myeloid leukemia. N Engl J Med. 2013;368:2059–74.

4. Figueroa ME, Lugthart S, Li Y, Erpelinck-Verschueren C, Deng X, Christos PJ, et al. DNA methylation signatures identify biologically distinct subtypes in acute myeloid leukemia. Cancer Cell. 2010;17:13–27.

5. Jones CL, Stevens BM, Pollyea DA, Culp-Hill R, Reisz JA, Nemkov T, et al. Nicotinamide Metabolism Mediates Resistance to Venetoclax in Relapsed Acute Myeloid Leukemia Stem Cells. Cell Stem Cell. 2020;27:748–64.e4.

6. Pollyea DA, Stevens BM, Jones CL, Winters A, Pei S, Minhajuddin M, et al. Venetoclax with azacitidine disrupts energy metabolism and targets leukemia stem cells in patients with acute myeloid leukemia. Nat Med. 2018;24:1859–66.

7. Laverdière I, Boileau M, Neumann AL, Frison H, Mitchell A, Ng SWK, et al. Leukemic stem cell signatures identify novel therapeutics targeting acute myeloid leukemia. Blood Cancer J. 2018;8:1–16.

8. Eppert K, Takenaka K, Lechman ER, Waldron L, Nilsson B, van Galen P, et al. Stem cell gene expression programs influence clinical outcome in human leukemia. Nat Med. 2011;17:1086–93.

9. Ng SWK, Mitchell A, Kennedy JA, Chen WC, McLeod J, Ibrahimova N, et al. A 17-gene stemness score for rapid determination of risk in acute leukaemia. Nature. 2016;540:433–7.

10. van Galen P, Hovestadt V, Wadsworth MH Ii, Hughes TK, Griffin GK, Battaglia S, et al. Single-Cell RNA-Seq Reveals AML Hierarchies Relevant to Disease Progression and Immunity. Cell. 2019;176:1265–81.e24.

11. Jung N, Dai B, Gentles AJ, Majeti R, Feinberg AP. An LSC epigenetic signature is largely mutation independent and implicates the HOXA cluster in AML pathogenesis. Nat Commun. 2015;6:8489.

12. van Rhenen A, Feller N, Kelder A, Westra AH, Rombouts E, Zweegman S, et al. High stem cell frequency in acute myeloid leukemia at diagnosis predicts high minimal residual disease and poor survival. Clin Cancer Res. 2005;11:6520–7.

13. Misaghian N, Ligresti G, Steelman LS, Bertrand FE, Bäsecke J, Libra M, et al. Targeting the leukemic stem cell: the Holy Grail of leukemia therapy. Leukemia. 2009;23:25–42.

14. Bonnet D, Dick JE. Human acute myeloid leukemia is organized as a hierarchy that originates from a primitive hematopoietic cell. Nat Med. 1997;3:730–7.

15. Shlush LI, Mitchell A, Heisler L, Abelson S, Ng SWK, Trotman-Grant A, et al. Tracing the origins of relapse in acute myeloid leukaemia to stem cells. Nature. 2017;547:104–8.

16. Corces MR, Buenrostro JD, Wu B, Greenside PG, Chan SM, Koenig JL, et al. Lineage-specific and single cell chromatin accessibility charts human hematopoiesis and leukemia evolution. Nat Genet. 2016;48:1193–203.

17. Duployez N, Marceau-Renaut A, Villenet C, Petit A, Rousseau A, Ng SWK, et al. The stem cell-associated gene expression signature allows risk stratification in pediatric acute myeloid leukemia. Leukemia. 2019;33:348–57.

18. Ng SW, Murphy T, King I, Zhang T, Mah M, Lu Z, et al. A Clinical Laboratory-Developed LSC17 Stemness Score Assay for Rapid Risk Assessment of Acute Myeloid Leukemia Patients. Blood Adv [Internet]. 2021; Available from: http://dx.doi.org/10.1182/bloodadvances.2021005741

19. Xie SZ, Kaufmann KB, Wang W, Chan-Seng-Yue M, Gan OI, Laurenti E, et al. Sphingosine-1-Phosphate Receptor 3 Potentiates Inflammatory Programs in Normal and Leukemia Stem Cells to Promote Differentiation. Blood Cancer Discovery. 2021;2:32–53.

20. van Galen P, Mbong N, Kreso A, Schoof EM, Wagenblast E, Ng SWK, et al. Integrated Stress Response Activity Marks Stem Cells in Normal Hematopoiesis and Leukemia. Cell Rep. 2018;25:1109–17.e5.

21. Paczulla AM, Rothfelder K, Raffel S, Konantz M, Steinbacher J, Wang H, et al. Absence of NKG2D ligands defines leukaemia stem cells and mediates their immune evasion. Nature. 2019;572:254–9.

22. Lechman ER, Gentner B, Ng SWK, Schoof EM, van Galen P, Kennedy JA, et al. miR-126 Regulates Distinct Self-Renewal Outcomes in Normal and Malignant Hematopoietic Stem Cells. Cancer Cell. 2016;29:214–28.

23. Jones CL, Stevens BM, D’Alessandro A, Reisz JA, Culp-Hill R, Nemkov T, et al. Inhibition of Amino Acid Metabolism Selectively Targets Human Leukemia Stem Cells. Cancer Cell. 2018;34:724–40.e4.

24. Chao MP, Takimoto CH, Feng DD, McKenna K, Gip P, Liu J, et al. Therapeutic Targeting of the Macrophage Immune Checkpoint CD47 in Myeloid Malignancies. Front Oncol [Internet]. 2020;9. Available from: https://www.frontiersin.org/articles/10.3389/fonc.2019.01380/full

25. Jin L, Hope KJ, Zhai Q, Smadja-Joffe F, Dick JE. Targeting of CD44 eradicates human acute myeloid leukemic stem cells. Nat Med. 2006;12:1167–74.

26. Jin L, Lee EM, Ramshaw HS, Busfield SJ, Peoppl AG, Wilkinson L, et al. Monoclonal antibody-mediated targeting of CD123, IL-3 receptor alpha chain, eliminates human acute myeloid leukemic stem cells. Cell Stem Cell. 2009;5:31–42.

27. Majeti R, Chao MP, Alizadeh AA, Pang WW, Jaiswal S, Gibbs KD, et al. CD47 is an adverse prognostic factor and therapeutic antibody target on human acute myeloid leukemia stem cells. Cell. 2009;138:286–99.

28. Pabst C, Krosl J, Fares I, Boucher G, Ruel R, Marinier A, et al. Identification of small molecules that support human leukemia stem cell activity ex vivo. Nat Methods. 2014;11:436–42.

29. Arnone M, Konantz M, Hanns P, Paczulla Stanger AM, Bertels S, Godavarthy PS, et al. Acute Myeloid Leukemia Stem Cells: The Challenges of Phenotypic Heterogeneity. Cancers [Internet]. 2020;12. Available from: http://dx.doi.org/10.3390/cancers12123742

30. Pollyea DA, Jordan CT. Therapeutic targeting of acute myeloid leukemia stem cells. Blood. 2017;129:1627–35.

31. Morita K, Wang F, Jahn K, Hu T, Tanaka T, Sasaki Y, et al. Clonal evolution of acute myeloid leukemia revealed by high-throughput single-cell genomics. Nat Commun. 2020;11:5327.

32. Schuringa JJ, Bonifer C. Dissecting Clonal Heterogeneity in AML. Cancer Cell. 2020;38:782–4.

33. Shlush LI, Zandi S, Mitchell A, Chen WC, Brandwein JM, Gupta V, et al. Identification of pre-leukemic hematopoietic stem cells in acute leukemia. Nature. 2014;506:328–33.

34. Anderson K, Lutz C, van Delft FW, Bateman CM, Guo Y, Colman SM, et al. Genetic variegation of clonal architecture and propagating cells in leukaemia. Nature. 2011;469:356–61.

35. Boer B de, Prick J, Pruis MG, Keane P, Imperato MR, Jaques J, et al. Prospective Isolation and Characterization of Genetically and Functionally Distinct AML Subclones. Cancer Cell. 2018;34:674–89.e8.

36. Ding L, Ley TJ, Larson DE, Miller CA, Koboldt DC, Welch JS, et al. Clonal evolution in relapsed acute myeloid leukaemia revealed by whole-genome sequencing. Nature. 2012;481:506–10.

37. Miles LA, Bowman RL, Merlinsky TR, Csete IS, Ooi AT, Durruthy-Durruthy R, et al. Single-cell mutation analysis of clonal evolution in myeloid malignancies. Nature. 2020;587:477–82.

38. Hope KJ, Jin L, Dick JE. Acute myeloid leukemia originates from a hierarchy of leukemic stem cell classes that differ in self-renewal capacity. Nat Immunol. 2004;5:738–43.

39. Kreso A, Dick JE. Evolution of the cancer stem cell model. Cell Stem Cell. 2014;14:275–91.

40. Kaufmann KB, Garcia-Prat L, Liu Q, Ng SWK, Takayanagi S-I, Mitchell A, et al. A stemness screen reveals C3orf54/INKA1 as a promoter of human leukemia stem cell latency. Blood. 2019;133:2198–211.

41. Notta F, Mullighan CG, Wang JCY, Poeppl A, Doulatov S, Phillips LA, et al. Evolution of human BCR-ABL1 lymphoblastic leukaemia-initiating cells. Nature. 2011;469:362–7.

42. Takayama N, Murison A, Takayanagi S-I, Arlidge C, Zhou S, Garcia-Prat L, et al. The Transition from Quiescent to Activated States in Human Hematopoietic Stem Cells Is Governed by Dynamic 3D Genome Reorganization. Cell Stem Cell. 2021;28:488–501.e10.

43. Wagenblast E, Azkanaz M, Smith SA, Shakib L, McLeod JL, Krivdova G, et al. Functional profiling of single CRISPR/Cas9-edited human long-term hematopoietic stem cells. Nat Commun. Nature Publishing Group; 2019;10:1–11.

44. Notta F, Zandi S, Takayama N, Dobson S, Gan OI, Wilson G, et al. Distinct routes of lineage development reshape the human blood hierarchy across ontogeny. Science. 2016;351:aab2116.

45. Chèneby J, Ménétrier Z, Mestdagh M, Rosnet T, Douida A, Rhalloussi W, et al. ReMap 2020: a database of regulatory regions from an integrative analysis of Human and Arabidopsis DNA-binding sequencing experiments. Nucleic Acids Res. 2020;48:D180–8.

46. Murphy T, Ng SWK, Zhang T, King I, Arruda A, Claudio JO, et al. Trial in progress: Feasibility and validation study of the LSC17 score in acute myeloid leukemia patients. Blood. American Society of Hematology; 2019;134:2682–2682.

47. Wagenblast E, Araújo J, Gan OI, Cutting SK, Murison A, Krivdova G, et al. Mapping the cellular origin and early evolution of leukemia in Down syndrome. Science [Internet]. 2021;373. Available from: http://dx.doi.org/10.1126/science.abf6202

48. Guo G, Luc S, Marco E, Lin T-W, Peng C, Kerenyi MA, et al. Mapping Cellular Hierarchy by Single-Cell Analysis of the Cell Surface Repertoire. Cell Stem Cell. 2013;13:492–505.

49. Boutzen H, Saland E, Larrue C, de Toni F, Gales L, Castelli FA, et al. Isocitrate dehydrogenase 1 mutations prime the all-trans retinoic acid myeloid differentiation pathway in acute myeloid leukemia. J Exp Med. 2016;213:483–97.

50. Epigenetic Therapies for Cancer | NEJM [Internet]. Available from: https://www.nejm.org/doi/full/10.1056/NEJMra1805035?query=TOC

51. Shih AH, Abdel-Wahab O, Patel JP, Levine RL. The role of mutations in epigenetic regulators in myeloid malignancies. Nat Rev Cancer. 2012;12:599–612.

52. Hua X, Zhao W, Pesatori AC, Consonni D, Caporaso NE, Zhang T, et al. Genetic and epigenetic intratumor heterogeneity impacts prognosis of lung adenocarcinoma. Nat Commun. 2020;11:2459.

53. Chan SM, Thomas D, Corces-Zimmerman MR, Xavy S, Rastogi S, Hong W-J, et al. Isocitrate dehydrogenase 1 and 2 mutations induce BCL-2 dependence in acute myeloid leukemia. Nat Med. 2015;21:178–84.

54. Tovy A, Reyes JM, Gundry MC, Brunetti L, Lee-Six H, Petljak M, et al. Tissue-Biased Expansion of DNMT3A-Mutant Clones in a Mosaic Individual Is Associated with Conserved Epigenetic Erosion. Cell Stem Cell. 2020;27:326–35.e4.

55. Figueroa ME, Abdel-Wahab O, Lu C, Ward PS, Patel J, Shih A, et al. Leukemic IDH1 and IDH2 Mutations Result in a Hypermethylation Phenotype, Disrupt TET2 Function, and Impair Hematopoietic Differentiation. Cancer Cell. 2010;18:553–67.

56. Losman J-A, Looper RE, Koivunen P, Lee S, Schneider RK, McMahon C, et al. (R)-2-hydroxyglutarate is sufficient to promote leukemogenesis and its effects are reversible. Science. 2013;339:1621–5.

57. van Galen P, Kreso A, Mbong N, Kent DG, Fitzmaurice T, Chambers JE, et al. The unfolded protein response governs integrity of the haematopoietic stem-cell pool during stress. Nature. 2014;510:268–72.

58. Dobin A, Davis CA, Schlesinger F, Drenkow J, Zaleski C, Jha S, et al. STAR: ultrafast universal RNA-seq aligner. Bioinformatics. 2013;29:15–21.

59. McInnes L, Healy J, Saul N, Großberger L. UMAP: Uniform Manifold Approximation and Projection. J Open Source Softw. The Open Journal; 2018;3:861.

60. Buenrostro JD, Giresi PG, Zaba LC, Chang HY, Greenleaf WJ. Transposition of native chromatin for fast and sensitive epigenomic profiling of open chromatin, DNA-binding proteins and nucleosome position. Nat Methods. 2013;10:1213–8.

61. Li H, Durbin R. Fast and accurate short read alignment with Burrows-Wheeler transform. Bioinformatics. 2009;25:1754–60.

62. ENCODE Project Consortium. An integrated encyclopedia of DNA elements in the human genome. Nature. 2012;489:57–74.

63. Zhang Y, Liu T, Meyer CA, Eeckhoute J, Johnson DS, Bernstein BE, et al. Model-based analysis of ChIP-Seq (MACS). Genome Biol. 2008;9:R137.

64. Schep AN, Wu B, Buenrostro JD, Greenleaf WJ. chromVAR: inferring transcription-factor-associated accessibility from single-cell epigenomic data. Nat Methods. 2017;14:975–8.

65. Korsunsky I, Millard N, Fan J, Slowikowski K, Zhang F, Wei K, et al. Fast, sensitive and accurate integration of single-cell data with Harmony. Nat Methods. 2019;16:1289–96.

66. Itoh K, Tezuka H, Sakoda H, Konno M, Nagata K, Uchiyama T, et al. Reproducible establishment of hemopoietic supportive stromal cell lines from murine bone marrow. Exp Hematol. 1989;17:145–53.

67. Brinkman EK, Chen T, Amendola M, van Steensel B. Easy quantitative assessment of genome editing by sequence trace decomposition. Nucleic Acids Res. 2014;42:e168.

68. Notta F, Chan-Seng-Yue M, Lemire M, Li Y, Wilson GW, Connor AA, et al. A renewed model of pancreatic cancer evolution based on genomic rearrangement patterns. Nature. 2016;538:378–82.

