## Supplementary figures and images for "A primary patient-derived model for investigating functional heterogeneity within the human Leukemic Stem Cell Compartment"

### Supplementary Figures

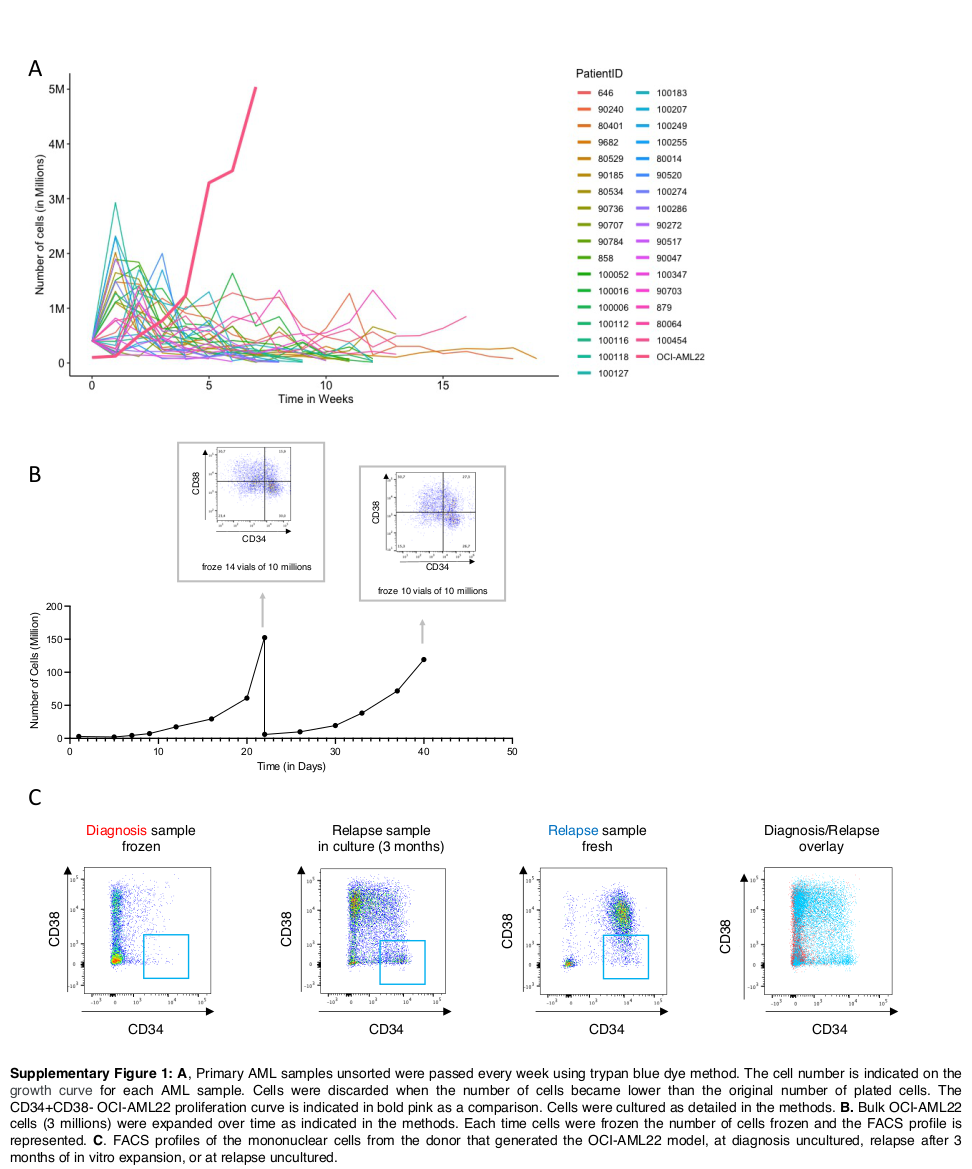
